# Characterization of the dual functions of *Leishmania* CK1.2 in both the parasite and the macrophage

**DOI:** 10.1101/2025.09.30.679250

**Authors:** Daniel Martel, Olivier Leclercq, Florent Dingli, Victor Laigle, Damarys Loew, Gerald F. Späth, Najma Rachidi

## Abstract

*Leishmania* CK1.2 (L-CK1.2) is a serine/threonine protein kinase essential for the survival of the protozoan parasite *Leishmania*, the causative agent of Leishmaniasis. This study investigates the dynamic localization pattern of L-CK1.2 and the broad spectrum of its interacting partners within the parasite. Using proteomic analysis and confocal microscopy, we identified 230 L-CK1.2-associated proteins across the parasite life stages, promastigotes (in the insect vector), and amastigotes, (in the phagolysosomes of host macrophages). This analysis revealed the ubiquitous presence of L-CK1.2 in various cellular structures, including the cytoskeleton, basal body, and flagellum. Using an *in vitro* system, sixty-four host L-CK1.2-associated proteins, involved in critical host biological processes such as immune response, apoptosis, and purine biosynthesis, were shown to interact with L-CK1.2. These processes are known to be regulated by *Leishmania* during infection. The study highlights the dual function of L-CK1.2, in the parasite (cis) and within the host cell (trans), positioning this kinase as a key player in host-pathogen interactions. This work provides a comprehensive map of L-CK1.2 interactions and suggest its potential importance in regulating intracellular *Leishmania* survival, providing potential therapeutic targets for Leishmaniasis. Furthermore, given the evolutionary conservation of CK1.2 across other parasitic organisms, our findings may have broader implications for understanding and managing parasitic infections.

## Introduction

CK1 family members are serine/threonine protein kinases ubiquitously expressed in eukaryotic organisms (1). They are involved in a wide range of important cellular processes, such as membrane trafficking or vesicular transport from yeast to humans (1). Due to its broad spectrum of action, CK1 activity and expression are tightly regulated by several mechanisms, including phosphorylation and subcellular sequestration (1). Indeed, members of the CK1 family are associated with many subcellular structures and organelles through interactions with specific partners ranging from circadian clock to cytoskeletal elements, (2–7), reflecting the strong interconnection of CK1 localisation and its functions. Defects in regulation, localisation or the introduction of mutations in the CK1 coding sequence are often associated with important diseases like cancer or neurodegenerative diseases (8) (9) (10). Regulation of CK1 pathways has also been linked to infectious diseases. For instance, inhibiting host CK1 suppresses yellow fever virus replication (11), highlighting the crucial role of CK1 in host-pathogen interactions. The protozoan parasite *Leishmania* is the causative agent of Leishmaniasis, a potentially life-threatening disease with at least three distinct clinical forms, cutaneous, mucocutaneous and the fatal visceral leishmaniasis. *Leishmania* has two developmental stages, an extracellular promastigote stage that proliferates inside the insect vector, and an intracellular amastigote stage that develops and multiplies inside the phagolysosomes of macrophages. The *Leishmania* genome encodes for six CK1 paralogs (12), of which only CK1.2 and CK1.4 are secreted by the parasite and were shown to be essential for parasite survival (13, 14). Several lines of evidence suggest that CK1.2 is released within the host cell as free protein or via extracellular vesicles (15–17). The adaptation of L-CK1.2 to two distinct cellular environments, driven by the high conservation of its protein sequence in *Leishmania* spp. and strong identity to its human orthologs, suggests that this protein has been shaped by selective pressures acting both on its functions within the host cell (12, 18). Indeed, our previous study identified potential substrates of L-CK1.2 in the macrophage, indicating its ability to regulate multiple host biological processes, such as ‘viral & symbiotic interaction’, ‘actin cytoskeleton organization’ and ‘apoptosis’ (19). This potentially dual functions, in *cis* in the parasite and in *trans* in the host macrophage (12, 18, 19), suggest *Leishmania* CK1.2 as a key player in host-pathogen interactions. Despite its importance in parasite biology and infection, the functions of L-CK1.2 remain poorly understood, beyond its known interactions with and phosphorylation of heat shock proteins within the parasite, and its ability to phosphorylate host targets such as IFNAR1, IFNAR2 (interferon-α/β receptors)(12, 13, 19–22). The essential nature of this kinase and its involvement in most cellular functions present major challenges for functional analysis using genetic approaches. To overcome these limitations and gain insights into the cis and trans functions of L-CK1.2, we chose a complementary approach determining and comparing its interactomes in both the parasite and the host cell. Our proteomic analysis unveils a repertoire of 230 interacting partners for L-CK1.2 across both amastigote and promastigote life stages of the parasite, with Gene Ontology enrichment analyses indicating its ubiquitous nature and multifaceted localization within various organelles. Applying confocal microscopy, we confirmed the localization of L-CK1.2 at these key subcellular sites, including the cytoskeleton, basal body, and flagellum. We used an *in vitro* approach to further identify 64 host proteins interacting with L-CK1.2, which again are implicated in a wide range of biological processes such as immune response, apoptosis and purine biosynthetic process. These biological processes are also modulated during *Leishmania* infection, strongly suggesting the involvement of L-CK1.2 in subverting host cell signalling. Our results suggest that certain pathways identified in the parasite as potential targets of L-CK1.2 are similarly found in the host. Collectively, our data provide additional evidence for the essential role of L-CK1.2, as a central component of the trans-signalling network between *Leishmania* and the macrophage. This dual biological function opens exciting avenues for future research on the co-evolution of parasite and macrophage biology, and the discovery of host-directed therapeutic interventions.

## Materials and Methods

### Leishmania cell lines

All the parasite cell lines used in this study were derived from *L. donovani* axenic 1S2D (MHOM/SD/62/1S-CL2D) clone *Ld*Bob, obtained from Steve Beverley, Washington University School of Medicine, St. Louis, MO. Promastigotes were cultured and differentiated into axenic amastigotes as described previously (23). Parasites cell lines were grown in media with 30 µg/mL hygromycin B (ThermoFisher Scientific Cat# 10687010) to maintain the pLEXSY-CK1.2-V5 or the empty pLEXSY plasmids. The transgenic *L. donovani* cell lines containing either the pLEXSY or pLEXSY-CK1.2-V5-HIS_6_ (pLEXSY-CK1.2-V5) vectors, corresponding to the mock or expressing *Leishmania major* CK1.2 tagged with V5 and HIS_6_, respectively, were described previously (18). L-CK1.2-V5 in this strain was shown to be active, functional, able to rescue a decrease in the activity of endogenous CK1.2 and its ectopic expression did not alter the parasite phenotypes.

### Immunofluorescence

To confirm the localisation of CK1.2 inferred from the interactome dataset, we used the *Leishmania donovani* mock and CK1.2-V5 parasites (see Mat & Met and (18)). Although, CK1.2-V5 is expressed ectopically, its abundant is twice as less as the endogenous CK1.2, precluding the risk of localisation artefact due to overexpression. Logarithmic phase promastigotes or axenic amastigotes (48h after shift at 37°C and pH5.5) were resuspended at 2x10^6^ parasites per mL in Dulbecco’s Phosphate Buffer Saline (DPBS) (Gibco) from at least three biological replicates, and 500 µL were added to poly-L-lysine-coated coverslips placed in a 24-well plate. Plates were centrifuged 10 min at 1200 g at room temperature to settle parasites onto the coverslips. For fixation alone, cells were washed three times with DPBS and fixed in 4% paraformaldehyde (PFA) in DPBS for 15 min at room temperature. For cytoskeleton preparation, the protocol was adapted from (24). Briefly, cells were washed three times with DPBS, treated with 0.125% Nonidet 40 (Fluka BioChemika Cat# 74385) in PIPES buffer (100 mM piperazine-N,N-bis(2-ethanesulfonic acid) (PIPES) pH6.8, 1 mM MgCl_2_) for 2 minutes at room temperature and washed twice for 5 minutes in PIPES buffer. Cells were fixed in 4% PFA in DPBS for 15 min at room temperature. After PFA fixation, cells were washed three times in DPBS, neutralised 10 min with NH_4_Cl (50 mM in DPBS), and washed again three times in DPBS. For the immuno-labelling of PFA-fixed cells or cytoskeleton preparations, the samples were blocked with 10% filtered heat-inactivated fetal calf serum (FCS) containing 0.5 mg.mL^-1^ saponin in DPBS for 30 min at room temperature and then washed for 5 min in DPBS. The cells were then incubated with primary antibodies diluted in DPBS with 0.5% Bovine Serum Albumin (BSA) and 0.5 mg.mL^-1^ saponin for 1h at room temperature. Three washes of 10 min were performed and the secondary antibody diluted in DPBS with 0.5% BSA and 0.5 mg.mL^-1^ saponin was added. After one hour incubation at room temperature in the dark, cells were washed twice for 10 min in DPBS with 0.5% BSA and 0.5 mg.mL^-1^ saponin, and then twice in DPBS. Parasites were incubated with 5 µg.mL^-1^ Hoechst 33342 in DPBS for 8 min in the dark, washed twice with DPBS, one time with distilled water, air-dried, then mounted with slides using SlowFade Gold Antifade Mountant (ThermoFisher Scientific Cat# S36937). For methanol fixation, logarithmic phase promastigotes were washed twice in DPBS and resuspended at 2x10^7^ parasites per mL. 10^6^ parasites were spread onto poly-L-lysine coated slides and allowed to settle for 30 min in a humid chamber. Parasites were then fixed in methanol at -20°C for 3 minutes and rehydrated for 10 min in DPBS at room temperature. For immuno-labelling of methanol-fixed parasites, samples were blocked with 10% filtered heat-inactivated FCS in DPBS for 15 min at room temperature and washed for 5 min in DPBS. Then the cells were treated similarly as those fixed by PFA. The antibodies used were: mouse IgG2a anti-V5 tag monoclonal antibody (Thermo Fisher Scientific Cat# R960-25, RRID:AB_2556564) diluted at 1/200 (in PFA and methanol fixed parasites) or at 1/300 (in cytoskeleton preparations); rabbit anti-V5 tag polyclonal antibody (Abcam Cat# ab9116, RRID:AB_307024) diluted at 1/400; rabbit anti-LdCentrin polyclonal antibody (kind gift from Hira L. Nakhasi) diluted at 1/2000 (25); mouse IgG1 anti-IFT172 monoclonal antibody diluted at 1/200 (26); mouse anti-PFR2 L8C4 clone antibody diluted at 1/10 (27); mouse L1C6 anti-TbNucleolus monoclonal antibody diluted at 1/100 (kind gift from Keith Gull (28)); mouse IgG1 anti-α-tubulin monoclonal DM1A antibody (Sigma-Aldrich Cat# T9026, RRID:AB_477593) diluted at 1/400; chicken anti-Hsp70 and chicken anti-Hsp90 antibodies diluted at 1/200 (29). IgG subclass-specific secondary antibodies coupled to different fluorochromes were used for double labelling: anti-mouse IgG (H+L) coupled to AlexaFluor488 (1/200 (PFA- or methanol-fixed parasites)) or 1/300 (cytoskeleton preparations), Thermo Fisher Scientific Cat# A-21202, RRID:AB_141607)); anti-mouse IgG2a coupled to Cy3 (1/600; Jackson ImmunoResearch Labs Cat# 115-165-206, RRID:AB_2338695); anti-mouse IgG1 coupled to AlexaFluor647 (1/600; Thermo Fisher Scientific Cat# A-21240, RRID:AB_2535809); anti-rabbit IgG (H+L) coupled to AlexaFluor488 (1/400; Thermo Fisher Scientific Cat# A-21206, RRID:AB_2535792); anti-mouse IgG (H+L) coupled to AlexaFluor594 (1/200; Thermo Fisher Scientific Cat# A-21203, RRID:AB_2535789); anti-mouse IgG2a coupled to AlexaFluor488 (1/300; Thermo Fisher Scientific Cat# A-21131, RRID:AB_2535771); anti-mouse IgG1 coupled to AlexaFluor594 (1/300; Thermo Fisher Scientific Cat# A-21125, RRID:AB_2535767); anti-chicken IgY coupled to AlexaFluor594 (1/200; Jackson ImmunoResearch Labs Cat# 703-586-155, RRID:AB_2340378) and anti-rabbit IgG (H+L) coupled to AlexaFluor594 (1/400; Thermo Fisher Scientific Cat# A-21207, RRID:AB_141637).

### Confocal microscopy

Images were visualised using a Leica SP5 HyD resonant scanner Matrix screener inverted microscope equipped with a HCX PL APO CS 63x, 1.4 NA oil objective (Leica, Wetzlar, Germany). Triple or quadruple immunofluorescence was imaged with Leica Application Suite AF software (LAS AF; Leica Application Suite X, RRID:SCR_013673) after excitation of the Hoechst 33342 dye with a diode at a wavelength of 405 nm (452/75 Emission Filter), excitation of the AlexaFluor488 with an argon laser at a wavelength of 488 nm (525/50 Emission Filter), excitation of AlexaFluor594 with a diode DPSS at a wavelength of 561 nm (634/77 Emission Filter), excitation of Cy3 with a diode DPSS at a wavelength of 561 nm (595/49 Emission Filter), and excitation of AlexaFluor647 with a helium-neon laser at a wavelength of 633 nm (706/107 Emission Filter). Images were scanned sequentially to minimise cross excitation between channels and each line was scanned twice and averaged to increase the signal-to-noise ratio. The pinhole aperture was set to 1 airy. Images were acquired with 8x zoom at a resolution of 1024×1024. Z-stacks were acquired at 0.082 µm intervals, deconvolved and rendered using either Fiji (RRID:SCR_002285) or Icy (RRID:SCR_010587) software (30) (http://icy.bioimageanalysis.org/).

### Deconvolution of z-stacks and chromatic aberration correction

All confocal images were processed and analysed by using the Huygens Professional software version 19.04 (Scientific Volume Imaging, Huygens Software, RRID:SCR_014237). Deconvolution of confocal z-stacks was optimised using the following settings: automatic estimation of the average background with the mode “Lowest” and area radius = 0.7, deconvolution algorithm CMLE, maximum number of iterations = 40, signal to noise ratio (SNR) = 20, quality change threshold = 0.05, iteration mode = optimised, brick layout = automatic. Theoretical point spread function (PSF) values were estimated for each *z*-stack. All deconvolved images were corrected for chromatic shifts and for rotational differences between different channels using the Chromatic Aberration Corrector (CAC) from Huygens Professional software (Scientific Volume Imaging, Huygens Software, RRID:SCR_014237). To calibrate the image corrections, multifluorescent 0.2 µm TetraSpeck microspheres (ThermoFisher Scientific Cat#T7280) mounted on SlowFade Gold Antifade mountant (ThermoFisher Scientific Cat# S36937) were imaged with identical acquisition parameters. Images were deconvolved similarly and were used to perform the chromatic aberration estimations with the cross-correlation method in CAC software. Corrections were saved as templates and applied for correction of the similarly acquired and deconvolved images in CAC.

### Co-localisation analysis of confocal images

Co-localisation analysis was performed with the Co-localisation Analyzer plug-in of the Huygens Professional software (Scientific Volume Imaging, Huygens Software, RRID:SCR_014237, v19.04). Processed cross-section images (deconvolved and corrected for chromatic aberrations) of the parasites were opened with this plug-in and Pearson correlation coefficients (PCC) were calculated for each parasite to quantify co-localisation. A PCC above 0.5 indicates co-localisation. Specific areas of the parasite were cropped from the whole image (basal body area, flagella pocket area, mitotic spindle and reduced mitotic spindle areas, flagellar pocket neck area and flagellar tip area) with the “crop” tool of the Huygens Professional software. Criteria to crop specific structures from the images were: (1) basal bodies - a region containing one or two dots representing the basal bodies in the channel CEN and excluding most of the flagellar pocket and flagellar pocket neck; (2) flagellar pocket area – a region stained with CK1.2 and CEN but excluding the basal bodies dots and the flagellum; (3) mitotic spindle – the entire Hoechst-unstained nucleus region in mitotic cells, containing the mitotic spindle in the TUB channel; (4) reduced mitotic spindle areas – only the regions containing the TUB staining in the mitotic nuclei, since CK1.2 is present in Hoechst-unstained regions where TUB staining is not visible, co-localisation of the entire Hoechst-unstained region is biased by a strong staining with CK1.2. We decided to crop further to analyse the colocalisation of CK1.2 and TUB in the mitotic spindle stained by TUB; (5) flagellar pocket neck – a region at the junction of the flagellum and the parasite body, excluding most of the flagellar pocket and basal bodies CK1.2 staining; and (6) flagellar tip – region at the tip of the flagellum, containing only a fraction of the rest of the flagellum. Rationale: since CK1.2 is present in Hoechst-unstained regions where TUB staining is not visible, co-localisation of the entire Hoechst-unstained region is biased by a strong staining with CK1.2. We decided to crop further to analyse the colocalisation of CK1.2 and TUB in the mitotic spindle stained by TUB. For the analysis of the colocalisation of CK1.2 and TUB in the mitotic spindle stained by TUB, we cropped further to restrict to the mitotic spindle as the co-localisation of the entire Hoechst-unstained region was biased by a strong staining with CK1.2. Pearson coefficients were calculated for these images. Pearson coefficients of the co-localisation in the basal body and flagellar pocket areas of (i) CK1.2-V5 with Centrin (CEN), IFT172, and DNA (Hoechst 33342, H); or (i) CEN with IFT172, from 14 images were plotted in scattered dot plots with the mean and standard deviation using GraphPad Prism 8.1.1 (GraphPad Software, GraphPad Prism, RRID:SCR_002798). Pearson coefficients of the co-localisation of CK1.2-V5 with tubulin in the mitotic spindle and reduced mitotic spindle areas from seven images were plotted similarly. Pearson coefficients of the co-localisation of CK1.2-V5 with Hsp90 from seven images and with Hsp70 from ten images were also plotted similarly.

### Epifluorescence microscopy and automated parasite detection

Images were visualised using a Zeiss upright widefield microscope equipped with Apotome2 grids and a Pln-Apo 63x, 1.4 NA oil objective (Zeiss). Light source used was a Mercury Lamp HXP 120, and following filters were used: DAPI (Excitation G365; dichroic FT 395; emission BP 420-470), FITC-A488-GFP (Excitation BP 455-495; dichroic FT 500; emission BP 505-555) and A594-TexasRed-mCherry-HcRed-mRFP (Excitation BP 542-582; dichroic FT 593; emission BP 604-644). Images were captured on an Axiocam MRm camera using ZEN Blue software. For comparison of different cell lines, identical parameters of acquisition were applied on all samples.

### Protein extraction, SDS-PAGE and Western blot analysis

Logarithmic phase promastigotes were washed in DPBS and protein extraction was performed as described previously (23). Ten micrograms of total protein were separated by SDS-PAGE, and transferred onto polyvinylidene difluoride (PVDF) membranes (Pierce). Membranes were blocked with 5% BSA in DPBS supplemented with 0.25% Tween20 (PBST) and incubated over night at 4°C with primary antibody mouse IgG2a anti-V5 tag monoclonal antibody (1/1000; Thermo Fisher Scientific Cat# R960-25, RRID:AB_2556564) in 2,5% BSA in PBST. Membranes were then washed in PBST and incubated with secondary antibody anti-mouse IgG (H+L) coupled to horseradish peroxidase (1/20000; ThermoFisher Scientific Cat# 32230, RRID:AB_1965958). Proteins were revealed by SuperSignal™ West Pico Chemiluminescent Substrate (ThermoFisher Scientific Cat# 34580) using the PXi image analysis system (Syngene) at various exposure times. Membranes were then stained with Bio-Safe Coomassie (Bio-Rad Cat #1610786) to serve as loading controls.

### Recombinant expression, purification of CK1.2-V5-His_6_ and protein kinase assay

*Escherichia coli* Rosetta (DE3) pLysS Competent Cells (Merck Cat# 70956) containing pBAD-thio-topo-L-CK1.2-V5-His_6_ and pBADthio-V5-His6 (control thioredoxin) were grown at 37°C and induced with arabinose (0,02% final) for 4h at room temperature (18). Cells were harvested by centrifugation at 10,000 g for 10 min at 4°C and the recombinant proteins were purified as described previously (18) (31). The eluates were supplemented with 15% glycerol and stored at −80°C.

### Production of macrophage extracts

Bone marrow derived macrophages (BMDM) were obtained from BALB/c ByJRj mice (32). Macrophages were lysed with RIPA buffer (150 mM NaCl, 1% Triton X-100, 20 mM Tris HCl [pH 7.4], 1% NP-40, 1 mM EDTA) supplemented with complete protease inhibitor cocktail (Roche Applied Science, IN) and with 1 mM sodium orthovanadate and 1 mM PMSF. Macrophage lysates were then vortexed and, after sonication, clarified by centrifugation.

### Immuno-precipitation

#### Leishmania IP

*L. donovani* logarithmic phase promastigotes or axenic amastigotes (48h post differentiation) containing either pLEXSY (mock) or pLEXSY-CK1.2-V5-His6 (expressing CK1.2-V5-His6) were collected and total protein were extracted, as described previously (23). Six (promastigotes) or twelve milligrams (axenic amastigotes) of total proteins were used at 2 mg/mL concentration in RIPA lysis buffer for the immuno-precipitation. The lysates were pre-cleared for 30 min with 1.5 mg of fresh magnetic beads coupled to protein G (ThermoFisher Scientific Cat# 10003D), washed with DPBS-tween buffer (DPBS with 0.02% Tween-20, adjusted at pH 7.4) and mixed with cross-linked anti-V5-coupled magnetic beads according to the manufacturer protocol. After incubation with pre-cleared lysates, beads were washed then the bound proteins were eluted with the glycine elution buffer for the PRO condition and with glycine elution buffer followed by the NuPAGE loading buffer (invitrogen) and heating at 95°C for Ax AMA condition. Elutions were kept at -80°C before mass spectrometry analysis.

#### Host IP

The rL-CK1.2-V5, the control thioredoxin and the empty vector were incubated overnight with 142 μg BMDM protein lysates. After preclearing, the samples were incubated with magnetic beads coupled to protein G and cross-linked to anti-V5 antibody. Following washing, the immuno-precipitated proteins were eluted first with glycine elution buffer (elution 1 and 2) then with one elution of NuPAGE loading buffer (elution 3). Elutions 1 and 3 were used for MS analysis and proteins from elution 2 were separated on SDS-PAGE and stained with SYPRO Ruby to assess the IP (Figure 5A).

### Size exclusion chromatography

Parasites containing either pLEXSY_HYG_V5_CK1.2 were harvested from logarithmic-phase cultures by centrifugation at 1,600 × g for 10 min and washed three times in D-PBS without CaCl_2_ and MgCl_2_ (Gibco). Cells were resuspended either in 1X RIPA lysis buffer (Cell Signaling Technology) containing 1 mM Phenylmethylsulfonyl fluoride, 50 U/ml Benzonase nuclease, Purity > 90% (VWR) and protease inhibitors (cOmplete Mini, EDTA-free Protease Inhibitor Cocktail Tablets; Roche). Cells suspensions were incubated on ice for 45 min, disrupted by sonication for 5 min (20s pulse, 20s off) in ice and clarified by centrifugation at 12,000 g for 10 min at 4°C. Total protein extracts were quantified using RC DC Protein Assay Kit II (Bio-Rad) then used immediately. SEC fractionations were performed at 4°C on Superose 6 increase 10/300 GL column (Cytiva) using a ÄKTApurifier FPLC system (GE Healthcare) (P-900 pumps; INV-907 valve; M-925 mixing valves; FR-902 flow Restrictor; FRC-950 fraction collector; UV-900 detector; CU-950 controller) piloted by UNICORN 5.10 software (GE Healthcare). Column was equilibrated with 2 column volumes (CV) of elution buffer (50 mM sodium phosphate; 150 mM NaCl; pH 7.2) at a flow rate of 0.35 ml/min. Then 6.9 milligrams of total protein extract were load in 500 µl sample loop, injected onto the column, and eluted with 1 CV of elution buffer at a flow rate of 0.35 ml/min. Eluted proteins were collected in 96 fractions of 200 µl. Fifteen microliters of each fraction were separated on a Novex NuPAGE 4 to 12% bis-Tris gel (ThermoFischer scientific) followed by Western blotting. Blots were probed with an anti-V5 antibody and an anti-mouse HRP-conjugated antibody. Signals were revealed using SuperSignal West Pico kit (Pierce), monitored with ChemiDoc imager (Bio-Rad) and quantified using Image Lab Software (Bio-Rad).

### Mass spectrometry analysis

MS analysis of the samples was done at the “Laboratoire de Spectrométrie de Masse Protéomique (LSMP)” platform at Institut Curie in Paris.

#### Leishmania IP

Proteins were run on SDS–PAGE gels (Invitrogen) without separation just as a clean-up step and stained with colloidal blue staining (LabSafe GEL Blue^TM^ G Biosciences). Gel slices were excised, and proteins were reduced with 10 mM DTT prior to alkylation with 55 mM iodoacetamide. In-gel digestion was performed using trypsin/Lys-C (Promega) overnight in 25 mM NH4HCO3 at 30 °C. Peptides were then extracted using 60/35/5 MeCN/H2O/HCOOH and vacuum concentrated to dryness. Samples were separated by chromatography using an RSLCnano system (Ultimate 3000, Thermo Scientific) coupled to an Orbitrap Fusion mass spectrometer (Q-OT-qIT, Thermo Fisher Scientific). We acquired Survey MS scans in the Orbitrap with the resolution set to a value of 120,000 and a 4 × 10^5^ ion count target. Each scan was recalibrated in real time by co-injecting an internal standard from ambient air into the C-trap. Tandem MS was performed by isolation at 1.6 Th with the quadrupole, HCD fragmentation with normalized collision energy of 35, and rapid scan MS analysis in the ion trap. The MS2 ion count target was set to 10^4^ and the max injection time was 100 ms. Only those precursors with charge state 2–7 were sampled for MS2. The dynamic exclusion duration was set to 60s with a 10 ppm tolerance around the selected precursor and its isotopes. The instrument was run in top speed mode with 3 sec cycles.

For identification, the data were searched against the Ld1S2D database (https://www.ncbi.nlm.nih.gov/bioproject/PRJNA396645, GCA_002243465.1) using Sequest HT through Proteome Discoverer (version 2.4). Enzyme specificity was set to trypsin and a maximum of two missed cleavages was allowed. Oxidized methionine and N-terminal acetylation were set as variable modifications and carbamidomethyl cysteine as fixed modification. The mass tolerances in MS and MS/MS were set to 10 ppm and 0.6 Da, respectively. The resulting files were further processed using myProMS v3.9.3 (https://github.com/bioinfo-pf-curie/myproms, (33)). False-discovery rate (FDR) was calculated using Percolator (34) and was set to 1% at the peptide level for the whole study. Label-free quantification was performed using peptide extracted ion chromatograms (XICs), re-extracted within conditions, and computed with MassChroQ v.2.2.21 (35). For protein quantification, XICs from proteotypic peptides shared between compared conditions (TopN matching) with missed cleavages were used. Median and scale normalization at peptide level was applied on the total signal to correct the XICs for each biological replicate (N=3 in both conditions). To estimate the significance of the change in protein abundance, a linear model (adjusted on peptides and biological replicates) was performed, and p-values were adjusted using the Benjamini–Hochberg FDR procedure. Proteins with at least two distinct peptides in two replicates of a same state, 2-fold enrichment and an adjusted p-value ≤ 0.05 were considered significantly enriched in sample comparisons. Proteins unique to a condition were also considered if they matched the peptides criteria. The mass spectrometry proteomics raw data have been deposited to the ProteomeXchange Consortium via the PRIDE (36) partner repository with the dataset identifier pxd033809.

#### Host IP

The elutions 1 and 3 were also run on SDS-PAGE without separation and only one gel slice containing the entire elution fraction was excised and used for nanoscale liquid chromatography coupled to tandem mass spectrometry (nanoLC-MS/MS) as described in the paragraph *Leishmania IP*. For protein identification, the data were searched against the databases containing SwissProt Mus Musculus (16745 sequences), contaminants and Uniprot *L. donovani* CK1.2 by using Sequest HT from Proteome Discoverer (version 2.4 or 2.1). The mass spectrometry proteomics raw data have been deposited to the ProteomeXchange Consortium via the PRIDE (36) partner repository with the dataset identifier: Project accession: PXD064006.

### Bioinformatics

The gene ontology analyses were performed with PANTHER (http://www.pantherdb.org/) (37) using *L. major* orthologs as input, or manually annotated using TriTrypDB (https://tritrypdb.org/tritrypdb/app) (38). STRING database, used to visualize protein complexes among the L-CKAP and L-CKAPhost, were merged to the interaction networks of L-CKAP*host* using Cytoscape software (version 3.7.1, https://string-db.org/) (39). The localisation of *T. brucei* proteins was determined using TrypTag website (http://tryptag.org/) (40). The localisation of *Leishmania mexicana* proteins was determined using LeishTag (https://www.leishtag.org/). For motif determination, we used SMART (http://smart.embl-heidelberg.de/). The dataset was analysed for protein-protein interactions and visualized using the STRING plugin (string, https://string-db.org/(41)) of the Cytoscape software package (version 3.8.2, https://cytoscape.org/(42)). Each node represents a substrate, and each edge represents a protein-protein interaction. The text was improved using ChatGPT 3.5 with the prompt, rewrite or improve (version 3.5, https://openai.com/chatgpt).

### Quantification and statistical analysis

Statistical analyses were performed with GraphPad Prism 8.1.1 (GraphPad Software, GraphPad Prism, RRID:SCR_002798) using unpaired t test (parametric test). Graphs were drawn using the same software. All errors correspond to the 95% confidence interval. Statistically significant differences are indicated with three (p<0.01), four (p<0.001) or five asterisks (p<0.0001). The number of samples analysed for each experiment is indicated in figure legends.

### Resource table

| REAGENT or RESOURCE | SOURCE | IDENTIFIER |
| --- | --- | --- |
| Antibodies |  |  |
| Anti-V5 tag IgG2a mouse monoclonal | Thermo Fisher Scientific | Cat# R960-25, RRID:AB_2556564 |
| Anti-Centrin from *L. donovani* rabbit polyclonal | (25) | N/A |
| Anti-IFT172 from *T. brucei* IgG1 mouse monoclonal | (26) | N/A |
| Anti-V5 tag IgG rabbit polyclonal | Abcam | Cat# ab9116, RRID:AB_307024 |
| Anti-PFR2 from *T. brucei* mouse polyclonal (L8C4) | (27) | N/A |
| Anti-nucleolus IgG from *T. brucei* mouse monoclonal (L1C6) | Keith Gull, University of Oxford, UK | N/A |
| Anti-alpha-tubulin IgG1 mouse monoclonal DM1A | Sigma-Aldrich | Cat# T9026, RRID:AB_477593 |
| Anti-mouse IgG (H+L) coupled to AlexaFluor488 | Thermo Fisher Scientific | Cat# A-21202, RRID:AB_141607 |
| Anti-mouse IgG2a coupled to Cy3 | Jackson ImmunoResearch Labs | Cat# 115-165-206, RRID:AB_2338695 |
| Anti-mouse IgG1 coupled to AlexaFluor647 | Thermo Fisher Scientific | Cat# A-21240, RRID:AB_2535809 |
| Anti-rabbit IgG (H+L) coupled to AlexaFluor488 | Thermo Fisher Scientific | Cat# A-21206, RRID:AB_2535792 |
| Anti-mouse IgG (H+L) coupled to AlexaFluor594 | Thermo Fisher Scientific | Cat# A-21203, RRID:AB_2535789 |
| Anti-mouse IgG2a coupled to AlexaFluor488 | Thermo Fisher Scientific | Cat# A-21131, RRID:AB_2535771 |
| Anti-mouse IgG1 coupled to AlexaFluor594 | Thermo Fisher Scientific | Cat# A-21125, RRID:AB_2535767 |
| Anti-chicken IgY coupled to AlexaFluor594 | Jackson ImmunoResearch Labs | Cat# 703-586-155, RRID:AB_2340378 |
| Anti-rabbit IgG (H+L) coupled to AlexaFluor594 | Thermo Fisher Scientific | Cat# A-21207, RRID:AB_141637 |
| Anti-mouse IgG (H+L) coupled to horseradish peroxidase | Thermo Fisher Scientific | Cat# 32230, RRID:AB_1965958 |
| Bacterial Strains |  |  |
| *Escherichia coli* Rosetta (DE3) pLysS Competent Cells | Merck | Cat# 70956 |
| Experimental Models: Organisms/Cell lines |  |  |
| *Ld*Bob cells with pLEXSY vector (mock) | (18) | N/A |
| *Ld*Bob cells with pLEXSY-CK1.2-V5-His<sub>6</sub> vector, expressing *Lma*CK1.2-V5-His<sub>6</sub> | (18) | N/A |
| Chemicals, Peptides, and Recombinant Proteins |  |  |
| Nonidet P40 (NP40) | Fluka BioChemika | Cat# 74385 |
| Ammonium Chloride (NH<sub>4</sub>Cl) | VWR Chemicals | Cat# 21235.297 |
| Carbobenzoxy-Leu-Leuleucinal (MG132) | Sigma-Aldrich | Cat# M7449 |
| SlowFade Gold Anti-Fade | ThermoFisher Scientific | Cat# S36937 |
| Recombinant thio-*Lma*CK1.2-V5-His<sub>6</sub> | (18) | N/A |
| Recombinant MBP | Sigma-Aldrich | Cat# M1891 |
| Recombinant DNA |  |  |
| Plasmid: pLEXSY-hyg2 (HygR) | Jena Bioscience | Cat# EGE-232 |
| Plasmid: pLEXSY-CK1.2-V5-His<sub>6</sub> (HygR) | (18) | N/A |
| Software and Algorithms |  |  |
| Icy | Icy | RRID:SCR_010587 |
| GraphPad Prism | GraphPad | RRID:SCR_002798 |
| CytExpert Software | Beckman Coulter | RRID:SCR_017217, v2.2.0.97 |
| Fiji | Fiji | RRID:SCR_002285 |
| Huygens Professional | Scientific Volume Imaging | RRID:SCR_014237 |
| Leica Application Suite AF (LAS AF) | Leica | RRID:SCR_013673 |

## Results

### *Leishmania* CK1.2 displays a pleiotropic localisation and interacts with numerous proteins in promastigotes and in axenic amastigotes

To determine L-CK1.2 localisation, we used the transgenic *L. donovani* Bob (*Ld*Bob) parasites carrying either the empty pLEXSY vector (referred to as ‘mock’) or pLEXSY-CK1.2-V5-His_6_ (referred to as ‘CK1.2-V5’), (18), previously characterised and functionally validated (18). Both strains were fixed with paraformaldehyde (PFA). L-CK1.2 was detected as intense punctate staining in the cytoplasm and the flagellum of PFA-fixed promastigotes expressing L-CK1.2-V5 (Figure 1Aa) but not in the corresponding mock control, which only showed weak background fluorescence (Figure 1Ab). The sum of the fluorescence intensity in the body of CK1.2-V5 promastigotes was significantly higher than that of the mock parasites with a p-value below 0.0001 (Figure S1A), indicating that this punctate staining is specific to CK1.2-V5. Similar localisations were also observed for axenic amastigotes (Figure 1Ac). This pleiotropic localization aligns with the diverse functions attributed to this protein kinase family. However, in the absence of a signal peptide for specific targeting, CK1 family members may primarily depend on protein-protein interactions for their localization (17).

To assess whether L-CK1.2 engages in protein-protein interactions in *Leishmania* promastigotes, we subjected V5-tagged L-CK1.2-containing lysates to size-exclusion chromatography (SEC) and analysed the distribution of the kinase across the elution profile. As shown in Figure 1B (blue circles), L-CK1.2-V5 was recovered in fractions spanning from 8 mL to 20 mL. Apart from a strong enrichment observed in fraction between 12 and 18 mL, the kinase was detectable evenly throughout most of the fractions, indicating that the bulk of cellular L-CK1.2 is incorporated into large complexes rather than existing as a free monomer. Indeed, the distribution of the total proteins is wider and detected up to fraction 25 mL (Figure 1B, pink circles). Collectively, these data support the notion that *Leishmania* CK1.2 predominantly participates in multi-protein complexes, underscoring the functional relevance of protein-protein interactions for this kinase.

**FIGURE 1:**
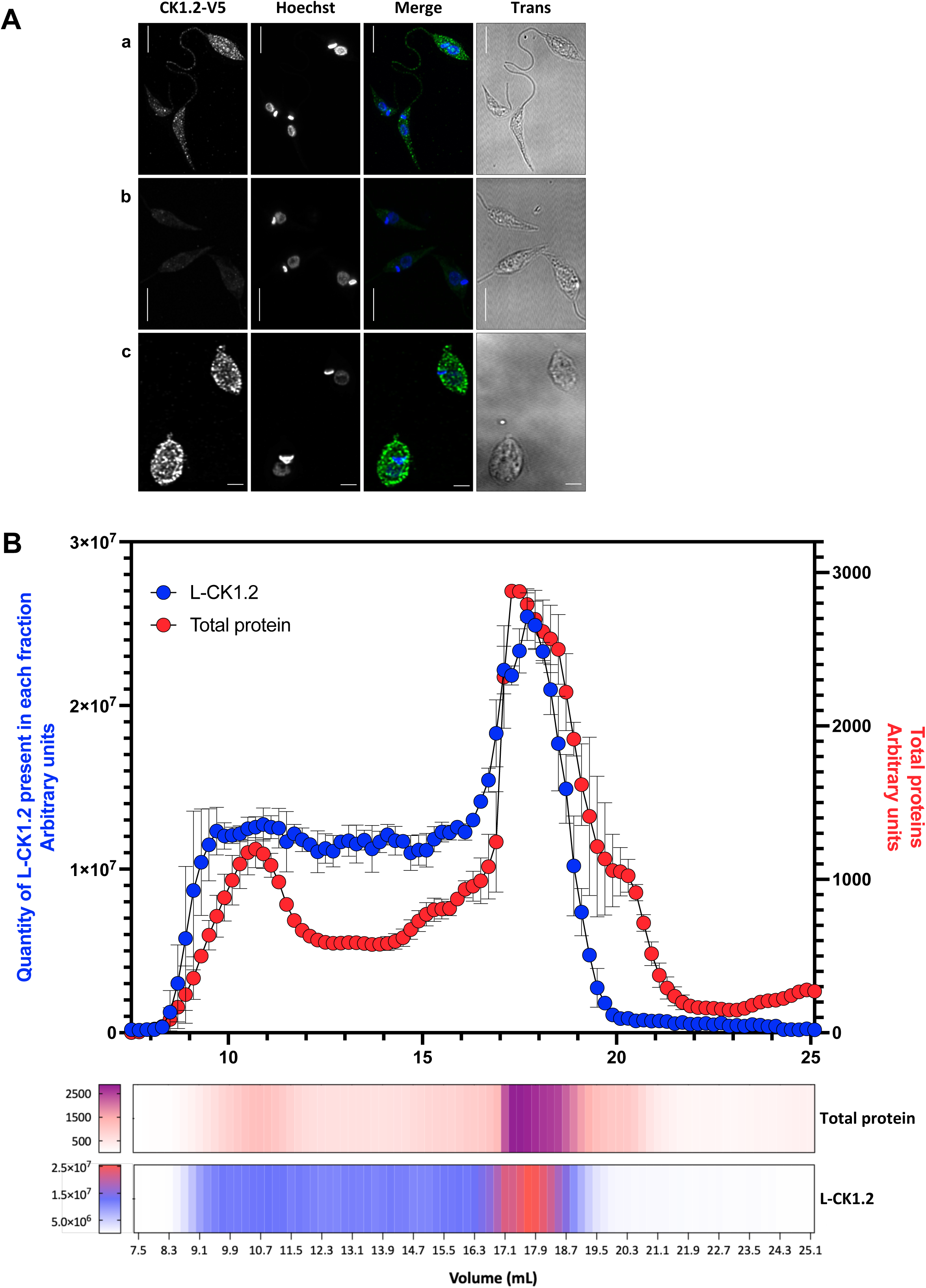
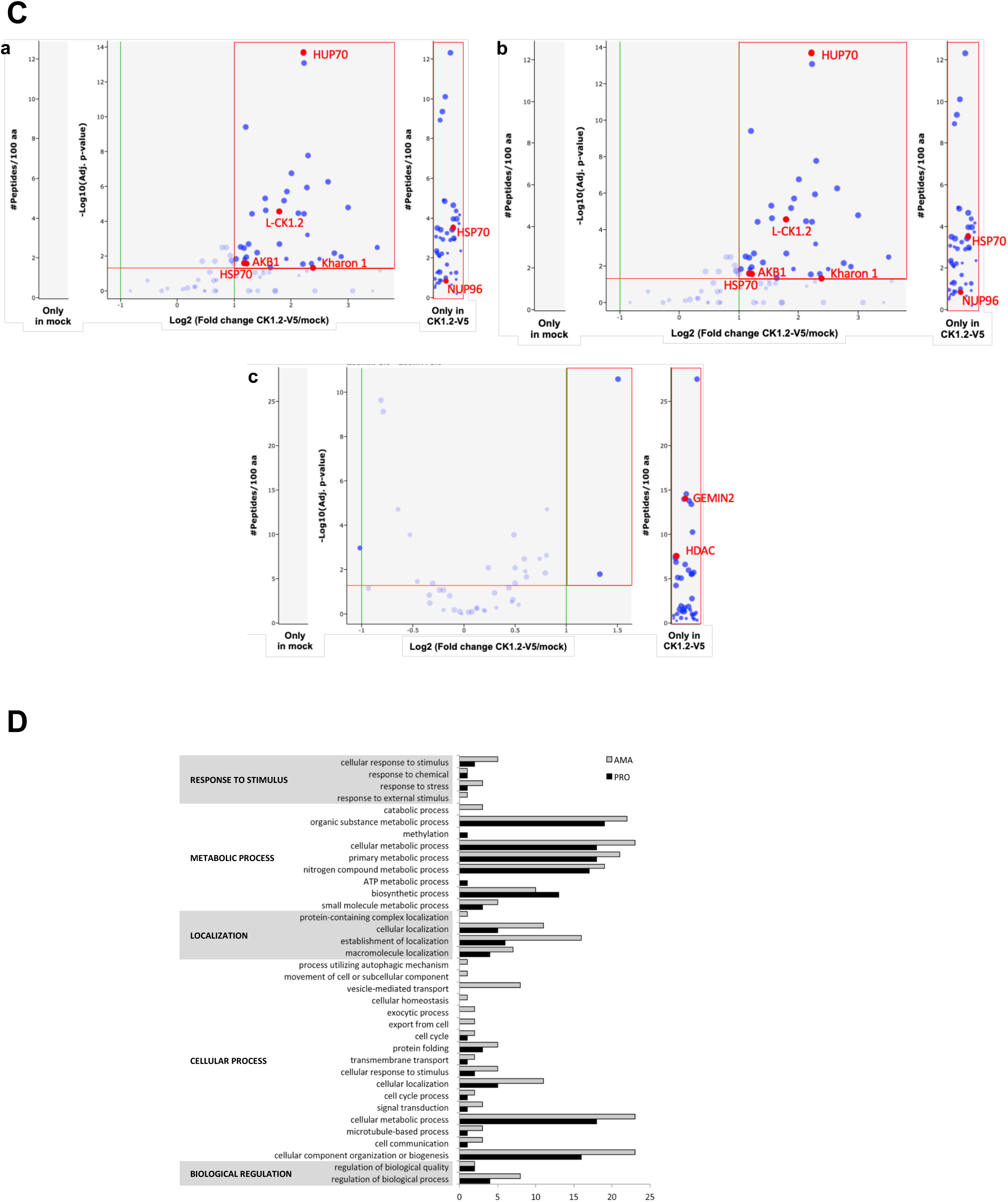
*Leishmania* CK1.2 displays a pleiotropic localisation and interacts with numerous proteins in promastigotes and in axenic amastigotes. (**A**) IFA of *Ld*Bob pLEXSY-CK1.2-V5 (a) and *Ld*Bob pLEXSY (mock, b) promastigotes, fixed with PFA. The confocal images show the anti-V5 staining (CK1.2-V5 or V5, green), Hoechst 33342 staining (H, blue), a merge and the transmission image (Trans). Scale bar, 2 µm. The pictures are maximum intensity projection of the confocal stacks containing the parasites. (c) IFA of *Ld*Bob pLEXSY-CK1.2-V5 axenic amastigotes, fixed with PFA. The single channel images show CK1.2-V5 (green) and Hoechst 33342 (H, blue) signals, the merge and the transmission image (Trans). Scale bar, 2 µm. The pictures are maximum intensity projection of the confocal stacks containing the parasites. Confocal stacks were deconvolved and corrected for chromatic aberration. The images presented in Figure 1A are representatives of at least 3 replicates. **(B)** Size-exclusion chromatography (SEC) of L-CK1.2-V5-containing complexes. 3.5 mg of protein, extracted in native conditions from *Ld*Bob pLEXSY-CK1.2-V5 (blue line) were separated by SEC on a superose 6 increase 10/300 GL columns. Fractions of 200 µl were collected and 15 µl of each fraction was separated by SDS-PAGE in denaturing conditions and analysed by Western blotting using an anti-V5 antibody (bottom panel). A blot for each strain, representative of the three replicates, is presented in the bottom panel. The quantification of the three replicates (dotted, dashed and plain lines) using the chemidoc imaging system (Biorad) is presented in the top panel. Each vertical line represents a fraction (from A1 to H12). MW= Molecular Weight. **(C)** Volcano plots representing the changes in protein abundance of LCKAPs in PRO (a), AxAMA elution E1 (b) and E2 (c). Proteins were selected if identified with at least 2 peptides in at least 2 replicates either only in the L-CK1.2-V5 conditions, or with a fold change above 2 and a p-value ≤ 0.05 compared to the mock conditions. Proteins highlighted in red are those with at least 2 distinct peptides in at least 2 replicates. **(D)** GO term enrichment for the ‘biological process’ in promastigotes and axenic amastigotes performed using PANTHER.db (http://www.pantherdb.org/, see also Table S4).

To identify the L-CK1.2 associated proteins (L-CKAP) influencing the regulation, functions or subcellular localisation of this kinase, we used mock or CK1.2-V5-expressing *Ld*Bob cell line (18). Three independent immuno-precipitations (IP) with logarithmic phase promastigotes (PRO) and axenic amastigotes (axAMA Elution1, E1 & Elution 2, E2) cell lysates were performed using a mouse monoclonal anti-V5 antibody. The eluted proteins were run on SDS-PAGE and analysed by quantitative label-free mass spectrometry. We identified 1081 proteins in promastigote and 516 in amastigote, from which respectively 645 (Figure 1Ca) and 341 (Figure 1Cb&c) were quantified (for the list of all quantified proteins, see Table S1). In addition to L-CK1.2 itself, 84 proteins in PRO and 146 in axAMA (Table S2) were selected as potential L-CKAPs, based on the following criteria: proteins identified with 2 or more distinct peptides, a fold change ratio CK1.2-V5/mock above 2 and a p-value below 0.05 in at least two replicates (Figure 1Ca-c, red square and Table S2), as well as those uniquely identified in CK1.2-V5 by at least 2 peptides or more in at least two replicates (Figure 1Ca-c, “only in CK1-V5” and Table S2). Forty-two L-CKAPs were shared between both parasite stages (Table S2, green). Half of the proteins identified in PRO and two third of those identified in AxAMA were stage-specific interactors of L-CK1.2 in our conditions. The dissimilarity between the two developmental stages becomes even more apparent when considering the enrichment of Gene Ontology (GO) terms associated with protein class (see Figure S1B and Table S3, PANTHER: http://www.pantherdb.org/) or biological processes (see Figure 1D, PANTHER: http://www.pantherdb.org/). In promastigotes, there is an over-representation of terms like ‘translational protein’ (Figure S1B) and an exclusive representation of ‘methylation’ and ‘ATP metabolic process’ (Figure 1D). In axenic amastigotes, terms such as ‘membrane traffic protein,’ ‘protein modifying enzyme,’ and ‘metabolite interconversion enzyme’ exhibit a higher representation (Figure S1B & Table S4) than in promastigotes, while terms such as ‘vesicle-mediated transport’, ‘exocytic process’ and ‘catabolic processes’ (Figure 1D) are exclusively associated with axenic amastigotes. These findings shed light on the potentially distinct functional characteristics exhibited by L-CK1.2 in the two stages.

### L-CK1.2 localisation in *Leishmania* parasites is consistent with that of its interaction partners

As the important cytoplasmic staining of L-CK1.2-V5 might mask other more specific localisations, promastigotes were treated with 0,125% NP-40 to permeabilise membranes and facilitate the release of the cytoplasmic pool of L-CK1.2, prior to PFA fixation and staining. The treatment conditions were selected after optimisation steps performed to minimise the impact of the detergent on the nucleus and kinetoplast morphology. We used organelle-specific markers for co-localisation with L-CK1.2 and also to demonstrate the integrity of the organelle. We observed organelle-specific signals (i) adjacent to the kinetoplast (Figure 2Aa, CK1.2-V5 & merge, white arrowhead), (ii) in the flagellar pocket region and along the flagellum (Figure 2Aa, CK1.2-V5 & merge, yellow arrowhead and arrow, respectively), as well as (iii) in the Hoechst-unstained region of the nucleus (Figure 2Aa, CK1.2-V5 & merge, white arrow). Similar localisations were observed in axenic amastigotes (Figure 2Ab, CK1.2-V5 & merge). These signals were reproducibly observed in all the samples that were analysed, including those fixed with methanol (as an example: 3 min, Figure S2), suggesting that the observed localisation is not a consequence of fixation processes. The organelle-specific localisation of L-CK1.2 might be due to interactions with specific L-CKAPs. To determine the potential localisation of these interacting partners in an exhaustive manner, we took advantage of the TrypTag (http://tryptag.org/ (40)), a community database providing the localisation of most proteins encoded in the genome of *Trypanosoma brucei* and we assumed that the L-CKAPs would have similar localisations than their syntenic *T. brucei* orthologs. When possible, we confirmed the localisation with the LeishTag database (https://www.leishtag.org/?pageType=landing). A total of 143 L-CKAPs were assigned subcellular localisation using TrypTag (Table S2, column: *T. brucei* & localisation in *T. brucei* orthologs), and 20 using LeishTag (Table S2, column: localisation in *L. mexicana* paralog & *L. mexicana*). The remaining 24 proteins lacked localisation data in both databases, or had no ortholog in *T. brucei*. The Gene Ontology enrichment for Cellular Component of the L-CKAPs was then established using the *T. brucei* orthologs, which are better annotated and Revigo software (http://revigo.irb.hr/ (43)) of Tritrypdb. L-CKAPs are localised in multiples organelles, including the cytosol, the nucleolus, the basal body, the flagellum, the flagellar pocket, or the nuclear pore complex (Figure 2B & Table S5 (blue for PRO, red for AxAMA and green for both)). Next, we performed co-localisation studies between L-CK1.2 and marker proteins specific for these organelles to validate its pleiotropic localization.

**FIGURE 2:**
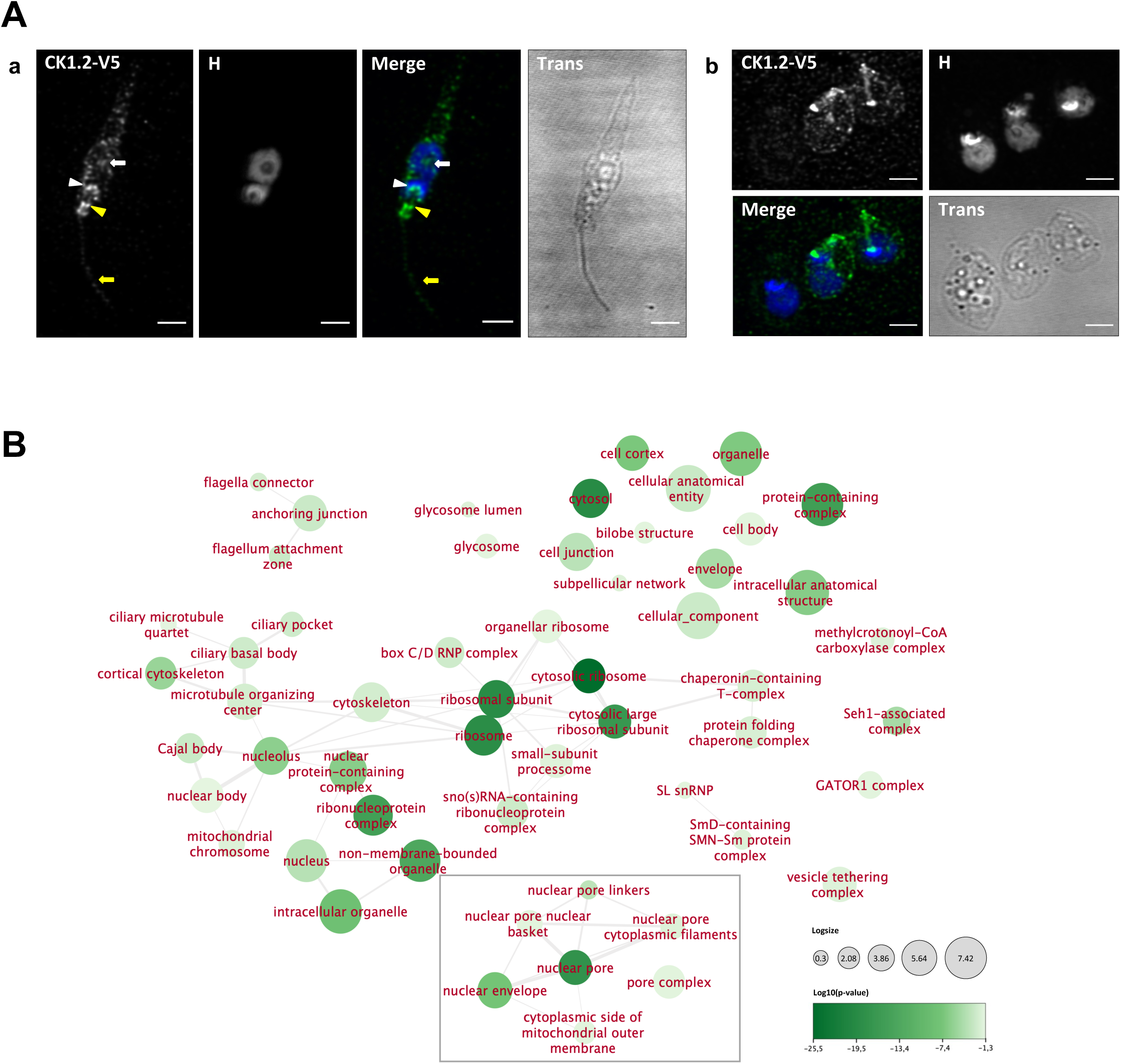
L-CK1.2 localisation in *Leishmania* parasites is consistent with that of its interaction partners. **(A)** IFA of *Ld*Bob pLEXSY-CK1.2-V5 promastigotes (a) and axenic amastigotes (b) obtained after detergent treatment followed by PFA fixation. Similar to (Figure 1A), except that the images are sum intensity projection of the confocal stacks containing the parasites. The images presented in Figure 2A are representatives of at least 3 replicates. **(B)** Graphical representation of the GO term enrichment for cellular component from the *Trypanosoma brucei* orthologs of both PRO and AxAMA interacting proteins using cytoscape app, Revigo (43) and tritrypdb GO enrichment (https://tritrypdb.org). The dataset of *Trypanosoma brucei* TREU927 has been used. The color of the node represents the (Log10(p-value), between value from - 25.55 to - 1.3). The size of the nodes represents the log size, which is the Log10(Number of annotations for GO Term ID in selected species in the EBI GOA database). Three percent of the strongest GO term pairwise similarities are designated as edges (grey) in the graph (43). The grey square highlights the nuclear pore complex.

### CK1.2 is localised in the granular zone of the nucleolus and redistributed to the mitotic spindle during mitosis

Forty-eight L-CKAPs have a potential localisation to the nucleolus (Table S2 and S5). Most of L-CK1.2 interacting partners in the nucleolus are involved in translation, with a wide range of 40S and 60S ribosomal proteins, as well as NOP56 (44). L-CK1.2 is also detected in the nucleolus as judged by Figure 2Aa (CK1.2-V5, white arrow), where CK1.2-V5 is detected in a Hoechst-unstained, sub-nuclear location. To confirm this result, the localisation of CK1.2-V5 in detergent-treated promastigotes was compared to that of L1C6 antibody, which recognises an unknown nucleolar protein (45). As expected, the L1C6-targeted antigen was detected in the centre of the nucleolus corresponding to the dense fibrillar zone involved in rDNA transcription (Figure 3A, merged image, red staining) (46), confirming the structural integrity of the nucleolus. In contrast, CK1.2-V5 was detected at the periphery of the nucleolus, as dotted staining around L1C6 (Figure 3A, merged image, green staining). This region corresponds to the granular component of the nucleolus, involved in the last steps of rRNA processing and ribosome biogenesis. Our results indicate that CK1.2 might be involved in rRNA processing rather than in rDNA transcription, which is consistent with its interaction with 40S and 60S ribosomal proteins, as well as the NOP56, a nucleolar protein involved in ribosome biogenesis, which orchestrates the correct assembly and the functioning of the box C/D small nucleolar ribonucleoproteins (snoRNPs) (44, 47–49). In addition to NOP56, L-CK1.2 interacts with members of NOP56 complex such as Fibrillarin (box C/D snoRNPs), Hel67 (LSU assembly factors), 60S ribosomal L7 protein, GTP-binding nuclear protein rtb2 and potentially the heat shock protein DNAJ (Figure 3B, (47)). Our interactome provides additional evidence supporting the localization of L-CK1.2 to the nucleolus and suggests a role for L-CK1.2 in ribosome processing.

**FIGURE 3:**
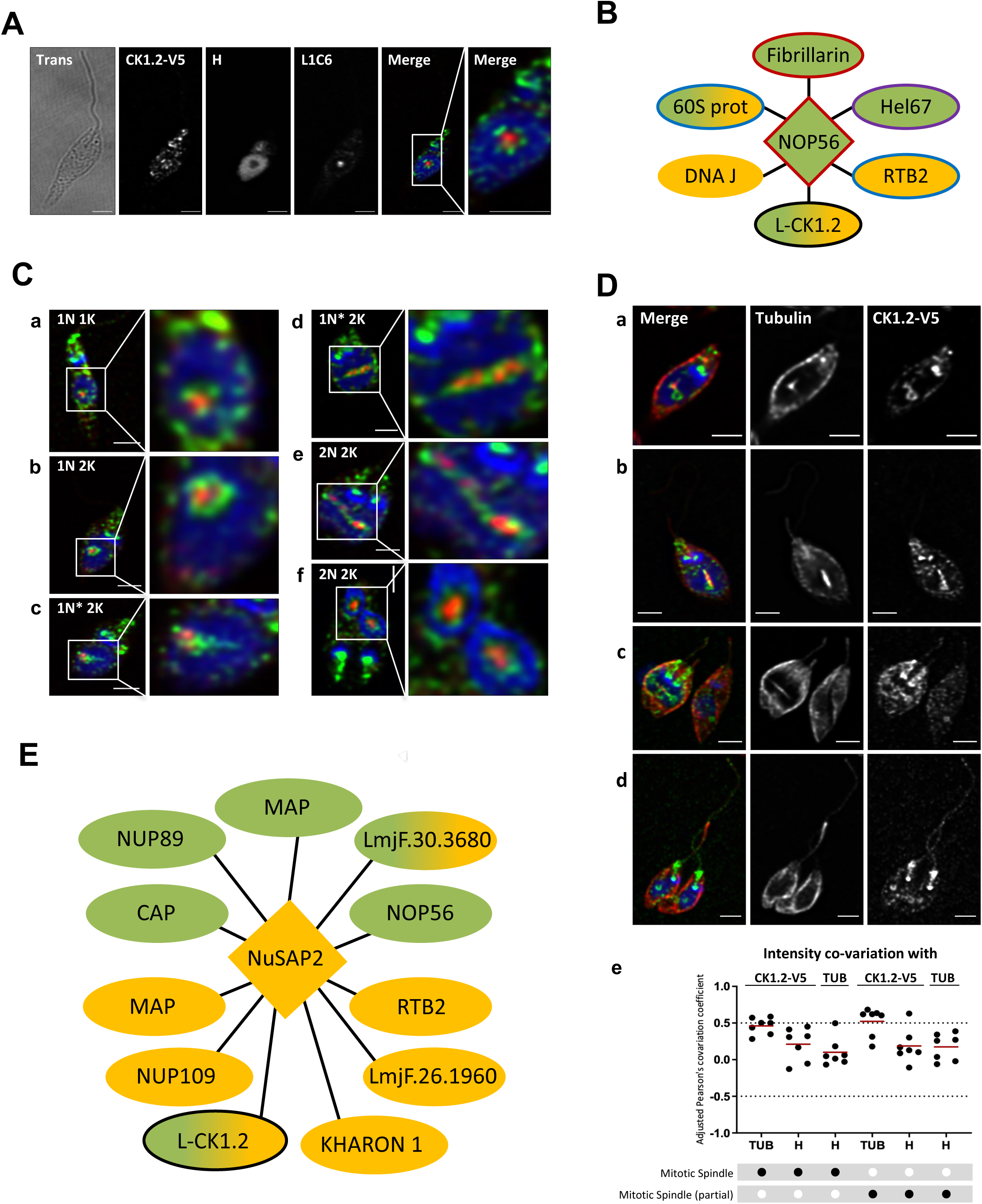
CK1.2 localises in the nucleolus and to the mitotic spindle. **(A)** IFA pictures of *Ld*Bob pLEXSY-CK1.2-V5 promastigotes obtained after detergent treatment followed by PFA fixation. The images show the single channel images of the transmission image (Trans), CK1.2-V5, Hoechst 33342 (H, blue) and L1C6 signals (red) and merged image. The right panel show a magnification of the nucleus region. Scale bar, 2 µm. These pictures are single stacks extracted from deconvolved confocal stacks corrected for chromatic aberration. **(B)** Subunits of NOP56 complex implicated in ribosomal subunits biogenesis found in our dataset. Fill colors: Green, detected in promastigotes; Yellow, detected in axenic amastigotes; and Green/Yellow, present in both life stage. Border colors: Red, corresponds to box C/D snoRNPs; Purple, corresponds to LSU assembly factors; Blue, putative ribosome assembly factors; and Black, potential interaction found in this study. **(C)** IFA pictures of *Ld*Bob pLEXSY-CK1.2-V5 promastigotes obtained after detergent treatment followed by PFA fixation and stained with anti-V5 (CK1.2-V5) and anti-L1C6 (nucleolus, L1C6) antibodies. Confocal images representing sequential events of mitosis revealed different localisation patterns of L1C6 nucleolar marker and CK1.2-V5. (a – f) The images show the merged image containing CK1.2-V5 (green), Hoechst 33342 (H) (blue) and L1C6 (red) signals and a magnification of the nuclear region. N=nucleus, K=kinetoplast. Scale bar, 2 µm or 1 µm for magnified images. These pictures are single stacks extracted from deconvolved confocal stacks corrected for chromatic aberration. See also Figure S3A. **(D)** IFA pictures of *Ld*Bob pLEXSY-CK1.2-V5 promastigotes obtained after detergent treatment followed by PFA fixation and stained with anti-V5 and anti-α-tubulin antibodies. Sequential images of various stages of cell division (a – d) showing the single channel images for α-tubulin signals and the merged images showing CK1.2-V5 (green), H (blue) and α-tubulin (red) signals. Scale bar, 2 µm. These pictures are single stacks extracted from deconvolved confocal stacks corrected for chromatic aberration. See also Figure S3B. (e) Dot plots showing Pearson’s covariation coefficients for different combination of signals in the entire mitotic spindle region and in a reduced mitotic spindle region containing also CK1.2-V5 signal. Regions of the spindle containing both CK1.2-V5 and tubulin were selected to evaluate whether L-CK1.2 localises to the spindle using Pearson’s covariation coefficients. For both regions, CK1.2-V5 signal was compared with α-tubulin (TUB) or Hoechst 33342 (H). The signal of TUB was also compared with H. Pearson’s covariation coefficients were measured from n=7 different confocal stacks which were deconvolved and corrected for chromatic aberration with Huygens Professional software. The plot was generated with GraphPad Prism software and the mean values are represented with red bold segments. **(E)** Subunits of NuSAP2 complex, found in our dataset, implicated in chromosome segregation. Fill colors: Green, detected in promastigotes; Yellow, detected in axenic amastigotes; and Green/Yellow, present in both life stage. Border colors: Black, potential interaction found in this study. The images presented in Figure 3 are representatives of at least 3 replicates.

The immunofluorescence experiment also unveils an additional function for nucleolar L-CK1.2. In dividing cells, in contrast to the typical segregation pattern observed for nucleolar components exemplified by L1C6 antigen (Figure 3C, red staining, panels a-f, and Figure S3A (50)), CK1.2 staining elongates from a wheel-shaped to a bar-shaped configuration that spans the entire length of the nucleus, following nucleolar elongation (Figure 3C, green staining, panels a-f, and Figure S3A). This localization pattern bears a resemblance to that of the mitotic spindle, and we have evidence suggesting an interaction between L-CK1.2 and α-tubulin, a component of the microtubules (Table S2). To determine whether L-CK1.2 is localised to the spindle during chromosome segregation, we asked whether L-CK1.2 co-localises with tubulin, a component of the spindle. As shown in Figure 3D (merged panels a-d) and Figure S3B, CK1.2-V5 co-localises with tubulin and thus with the mitotic spindle in specific areas, an observation supported by the mPc at or above 0.5 (Figure 3De, mitotic spindle).

During anaphase, CK1.2-V5 accumulates at each end of the elongated mitotic spindle (Figure 3Dc-merge panel, white arrows; Figure S3Bd, white arrows), similarly to the twinfilin-like protein (51). These findings suggest that *Leishmania* CK1.2 might play a role in regulating chromosome segregation similarly to TbCK1.2 (51, 52). Consistent with this hypothesis, Nucleus and spindle associated protein 2 (LdNuSAP2), a highly divergent ortholog of Saccharomyces ASE1/HumanPRC1/ArabidopsisMAP65 protein family that mediates spindle elongation, was identified in L-CK1.2 interactome (53) alongside eleven interacting partners of NuSAP2, suggesting that L-CK1.2 might be recruited to the spindle as part of the NuSAP2 complex (Figure 3E, (53)). Further investigations will be required to determine the mechanisms by which CK1.2 regulates mitosis.

A closer examination at potential complexes highlighted the specific association of CK1.2 with the nuclear pore complex subunits in a life-stage specific manner (NPC, Figure 2B, grey square). While L-CK1.2 interacts with only four NPC members in promastigotes (Figure S4C (blue squares) and Table S5), it interacts with eighteen NPC subunits in axenic amastigotes (Figure S4C (blue and red squares) and Table S5), including three hypothetical proteins. These proteins, LdBPK_091010.1, LdBPK_100580.1, LdBPK_281740.1, might be novel components of the NPC, as judged by the localization of their orthologs in *T. brucei* (40). These L-CKAPs cover most subcomplexes of the NPC but are particularly enriched in the outer ring (Figure S4C), suggesting a tight, life-stage specific interaction between L-CK1.2 and the nuclear pore in axenic amastigotes. Similarly to L-CK1.2, in *Trypanosoma brucei*, CK1.2 interacts with TbNUP158, a member of the outer ring (54) and Hrr25, CK1 ortholog in *S. cerevisiae*, interacts with and phosphorylates NUP53(55).

### CK1.2 localises to the basal body and the flagellum

Based on GO enrichment analysis of cellular component (Figure 2B), 12 L-CKAPs (Table S5) are predicted to locate to (i) the basal body that regulates flagellum duplication, (ii) the tripartite attachment complex (TAC) that links the segregation of the kinetoplasts to that of the basal bodies, or (iii) the transition fibers (TF), critical for flagellum assembly. Most of these interacting proteins were identified either in axenic amastigotes or in both AxAMA and PRO, none were specific of PRO. Several of these L-CKAPs are linked to the regulation of cell division and/or cytokinesis, such as *Leishmania* AKB1 or KHAP2 (13, 56, 57), while other are members of the Tripartite Attachment Complex (TAC) such as TAC60 (58). To determine whether L-CK1.2 localizes to these organelles, we selected two markers, centrin-4 to investigate the potential localisation of CK1.2-V5 to the basal body (Figure 4A, CEN, white arrow) and to the bilobe structure (Figure 4A, CEN, yellow arrow), and IFT172 to investigate the potential localisation of CK1.2-V5 to the transition fibers (Figure 4A, IFT172) (59–61). As shown in Figure 4A panel a, CK1.2-V5 does not co-localise neither with the kinetoplast DNA, as shown by the mean Pearson correlation coefficient below 0.5 (mPc=0.27 ± 0.164, Figure 4B), nor with the bilobed structure (Figure 4A panel a, yellow arrow and mPc of 0.47 ± 0.074, Figure 4B,). Instead, it co-localises with centrin-4 to the basal bodies (Figure 4A panel a, white arrows and mPc of 0.741 ± 0.050, Figure 4B), and with IFT172 to the transition fibers, as judged by Figure 4A panel b, white arrow and confirmed by an mPc of 0.805 ± 0.06 (Figure 4B), suggesting that L-CK1.2 might regulate flagellum duplication as well as be loaded onto the flagellum. Consistent with this finding, we confirmed the localisation of CK1.2-V5 to the axoneme (IFT172, Figure 4C, Merge and 3D-view), but not to the paraflagellar rod (PFR2, Figure 4D, panel 3). Noticeably, the abundance of L-CK1.2 in the flagellum is lower than that in the body of the parasite, which is consistent with the filtering existing at the entrance of the flagellum (62). The flagellar localisation of L-CK1.2, which aligns with published proteomic data from *T. brucei* and *Leishmania mexicana* flagellum (63, 64), as well as the identification of 19 flagellar L-CKAP (Table S5, section flagellum, (64)) indicates a potential function for L-CK1.2 in the flagellum. Additionally, L-CK1.2 phosphorylates HSP90 at Ser-289; blocking this modification, by introducing an S289A mutation, produces mutant parasites whose flagella are shortened by roughly 50 % (20). This finding suggests a regulatory role of L-CK1.2 in flagellar length.

**FIGURE 4:**
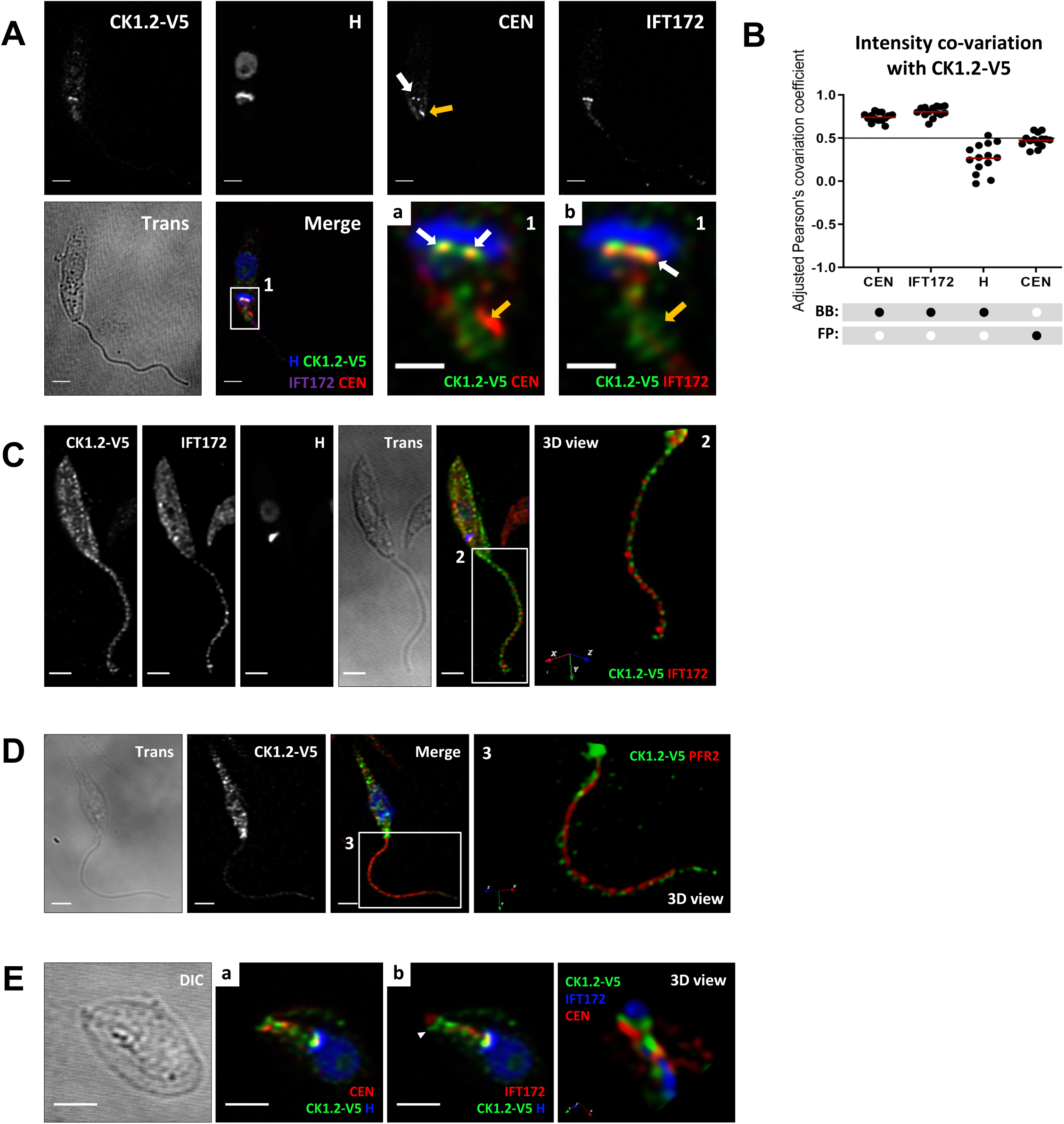
CK1.2 is localised to the basal bodies and the flagellum. **(A)** IFA of *Ld*Bob pLEXSY-CK1.2-V5 promastigotes obtained after detergent treatment followed by PFA fixation. Single channel images of the CK1.2-V5, Hoechst 33342 (H), centrin 4 (CEN) or IFT172 signals and the transmission image (Trans). The white arrow highlights the basal bodies and the yellow arrow the bilobe region. The merge panel shows CK1.2-V5 signal (green) merged with H (Hoechst 33342, blue), CEN (centrin 4, red) and IFT172 (cyan). The two right panels show a magnification of region 1 with H signal (blue) merged with (a) CK1.2-V5 (green) and CEN (red) and (b) IFT172 (red) and CK1.2-V5 (green) signals. Scale bar, 2 µm or 1 µm for magnified images. These pictures are single stacks extracted from deconvolved confocal stacks corrected for chromatic aberration. The white arrows highlight the basal bodies and the yellow arrow the bilobe region. **(B)** Dot plots showing Pearson’s covariation coefficients in the basal body (BB) or the flagellar pocket (FP) regions for different combination of signals. Pearson’s covariation coefficients were measured from n=14 different confocal stacks which were deconvolved and corrected for chromatic aberration with Huygens Professional software. The plot was generated with GraphPad Prism software and the mean values are represented with red bold segments. **(C)** IFA of *Ld*Bob pLEXSY-CK1.2-V5 promastigotes fixed by PFA and stained with the anti-V5 and anti-IFT172 antibodies. The left panels display the single channel images of the CK1.2-V5, IFT172, Hoechst 33342 (H) signals, the transmission image (Trans), and a merge of CK1.2-V5 (green), IFT172 (red) and H (blue) signals. The right panel shows a 3D-reconstruction (3D view) of the flagellum (image 2), with CK1.2-V5 signal (green) merged with IFT172 (red). **(D)** IFA of *Ld*Bob pLEXSY-CK1.2-V5 promastigotes obtained after detergent treatment followed by PFA fixation and staining with the anti-V5 and anti-PFR2 antibodies. The following images show CK1.2-V5, PFR2, Hoechst 33342 (H) signals, the transmission image (Trans), and finally the merge of CK1.2-V5 (green), PFR2 (red) and H (blue) signals. (image 3) 3D-reconstruction (3D view) of the flagellum (white square in (merge)), showing CK1.2-V5 signal (green) merged with PFR2 (red). Scale bars, 2 µm or 1 µm for 2D images. Pictures in (E) and (D, left panels) are single stacks extracted from deconvolved confocal stacks corrected for chromatic aberration. All confocal stacks containing the parasite were used for pictures (D, image 2 and E image 3). **(E)** IFA of *Ld*Bob pLEXSY-CK1.2-V5 axenic amastigotes obtained after detergent treatment followed by PFA fixation. The single channel and the merge images show CK1.2-V5, Hoechst 33342 (H, blue), centrin 4 (CEN, red) or IFT172 (cyan) signals and the transmission image (Trans). Scale bar, 2 µm. These pictures are single stacks extracted from deconvolved confocal stacks corrected for chromatic aberration. Panel a shows a merged of CK1.2-V5, Hoechst 33342 (H, blue) and centrin 4 (CEN, red); Panel b shows CK1.2-V5, Hoechst 33342 (H, blue), and IFT172 (cyan) signals; 3D-reconstruction of the flagellar pocket region and its neck from image (Figure 5Fa and b), which has been rotated. All confocal z-stacks containing the parasite were used for the 3D view. The images presented in Figure 4 are representatives of at least 3 replicates.

Based on the literature and our findings, *Leishmania* promastigote, alongside *Chlamydomonas reinhardtii* and *Trypanosoma brucei*, is one of the few eukaryotes exhibiting flagellar localization of CK1 (65). In axenic amastigotes, however, CK1.2-V5 does not colocalise with IFT172, but instead localises around the flagellum (Figure 4E, panel b and 3D view). These findings suggest that the role of CK1.2 in the flagellum is more likely associated with motility or flagellum assembly than in sensing.

### *Leishmania* CK1.2 interacts with host proteins

Several lines of evidence suggest that the L-CK1.2 is exported from the parasite in extracellular vesicles (EVs) and can phosphorylate host substrates, thereby contributing to the parasite ability to subvert macrophage biology (15, 19, 22). Because L-CK1.2 is expressed at low levels in both promastigotes and axenic amastigotes(18), conventional immunoprecipitation (IP) of the kinase from infected macrophages would not yield sufficient material for a comprehensive interactome analysis. Furthermore, the amount of host-localised pool of L-CK1.2 that can be harvested from *Leishmania*-infected BMDMs is well under the threshold needed for a robust IP-MS analysis. To circumvent these limitations, we adopted an *in-vitro* pull-down strategy that has previously been validated for mapping host–pathogen protein interactions in *Mycobacterium* (*66*).

Recombinant L-CK1.2-V5, produced in *E. coli*, was incubated overnight with macrophage protein lysates and immunoprecipitated using an anti-V5 antibody (18, 66) to identify the host L-CK1.2-associated proteins (L-CKAP*host*) by mass spectrometry analysis. Two controls were added, an empty vector, and a control-thioredoxin (an empty plasmid expressing thioredoxin-V5), as recombinant L-CK1.2-V5 is fused to thioredoxin at the N-terminus to improve its solubility. Three independent immuno-precipitations were performed using a mouse monoclonal anti-V5 antibody. The proteins were run on SDS-PAGE (Figure 5Aa) and the binding partners revealed by quantitative label-free mass spectrometry (Figure 5Aa, b &c). We identified 64 L-CKAP*host* in at least two replicates with at least two distinct peptides detected (Table S6). L-CK1.2-V5 interacts with a wide range of host protein classes, including chaperones and transporters, similar to the L-CKAPs identified in *Leishmania* (Figure 5B). This finding suggests that L-CK1.2 may perform similar functions in both the parasite and its host cell. The GO analysis for Biological Process (Figure S5A) revealed a significant enrichment for the terms ‘purine biosynthetic process’, ‘glycolytic process’ (ATP-dependent 6-phosphofructokinase, Pfkl and Glyceraldehyde-3-phosphate dehydrogenase, Gapdh), ‘mitochondrial electron transport’ (mitochondrial ATP synthase subunit, Atp5f1a, Atp5f1c and Atp5po, Cytochrome b-c1 complex subunit 8) and ‘mitochondrial membrane respiratory chain’ (Ndufa1, Ndufa10). These findings collectively suggest a potential involvement of L-CK1.2 in reprogramming host cell energy metabolism, which indeed undergoes a gradual shift from an M2-like respiratory metabolic state to an M1-like glycolytic energy profile during *Leishmania* infection (67). We also identified an enrichment of GO terms associated with infection, such as “immune response activation,” “apoptotic cell death,” and “response to stress”, suggesting a role for L-CK1.2 in subverting key host pathways that enable the parasite to establish its niche and promote intracellular survival (Figure S5A, (68)).

**FIGURE 5:**
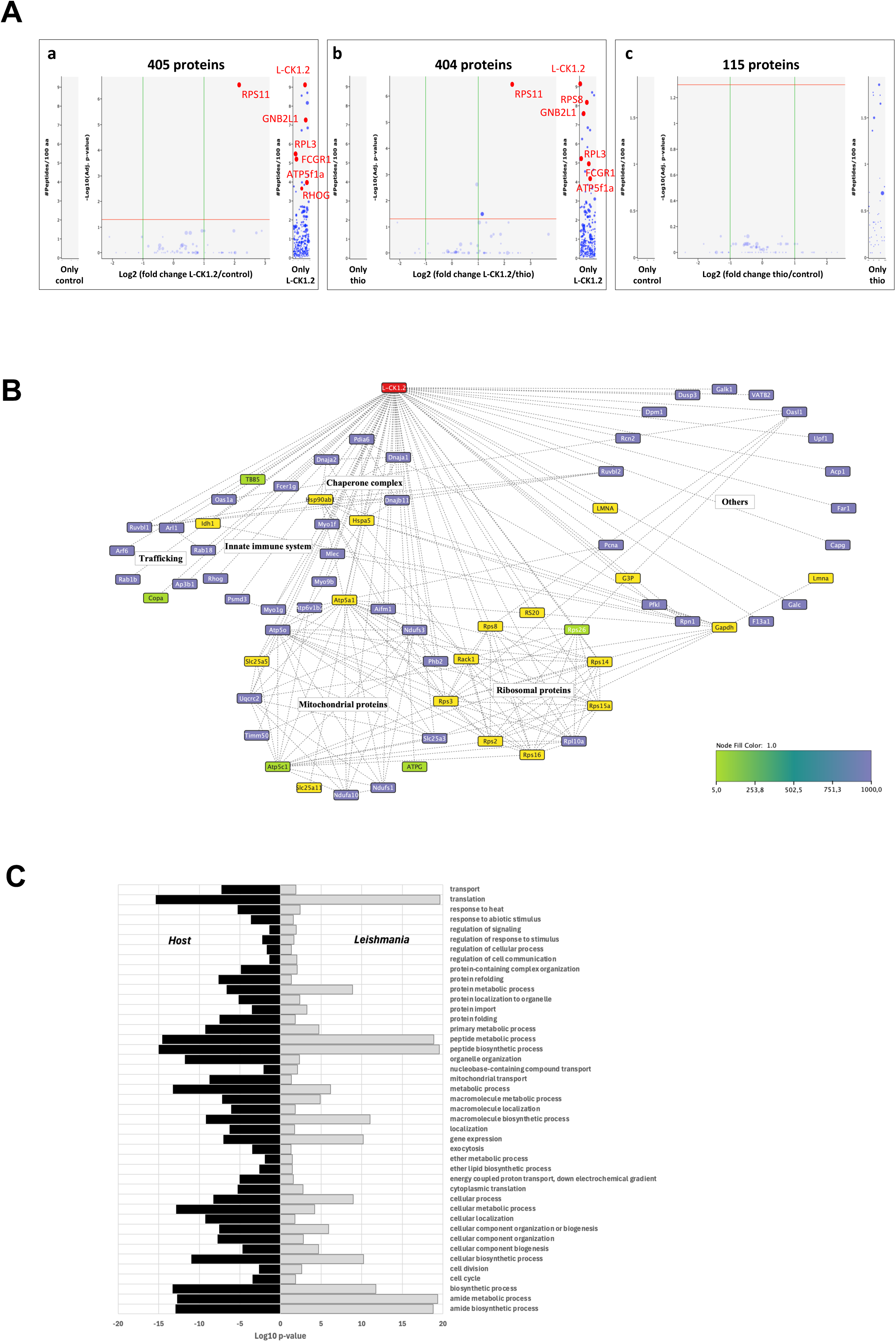
L-CK1.2 interacts with macrophage proteins. **(A)** The recombinant L-CK1.2-V5, the control-thioredoxin and the control were incubated overnight with BMDM protein lysates and immunoprecipitated using an anti-V5 antibody. The proteins were eluted by two successive elutions with glycine elution buffer (elution 1 and 2) followed by one elution with NuPAGE loading buffer (elution 3). Proteins from elutions 1 and 3 were used for MS analysis (Figure 5A a, b and c). Volcano plots representing the changes in protein abundance of L-CK1.2 vs control (a), of L-CK1.2 vs control thioredoxin (b) and of control thioredoxin vs control (c), including all the proteins identified with 1 peptide in at least 1 replicate. Proteins highlighted in red are those with at least 2 distinct peptides in at least 2 replicates. **(B)** Graphical representation of the affinity purification merged with the string app to visualise protein-protein interactions for host proteins using the cytoscape app. The classification in different categories is based on GO terms enrichment. The colour of the nodes represents the Fold change (see the legend on the figure). **(C)** GO term enrichment for the ‘biological process’ common between the host cell and *Leishmania donovani* performed using the cytoscape app (see also Table S7).

Notably, many of the biological processes enriched in the interactome are also present in the L-CK1.2-related host substratome we previously published (Figure S5B, Table S8 (19)). The most common biological processes shared between the host interactome and the host substratome are related to post-transcriptional regulation processes (Figure S5B, Table S8 (19)). Because of its potential dual role, L-CK1.2 has the ability to function in both the parasite and the macrophage. To determine whether L-CK1.2 targets the same pathways in both organisms, we analysed the intersection between biological processes associated with L-CK1.2 in the parasite and the host cell by combining the L-CKAP*host* and substratome datasets (this work and (19), Figure 5C, Table S7). Several biological processes were conserved in both organisms, representing the core of L-CK1.2 functions: (i) biosynthetic metabolic processes including RNA metabolic process and lipid biosynthetic process; (ii) translation; (iii) transport including endocytosis, and trafficking; (iv) response to stimuli, regulation of cell communication and signalling (Figure 5C). Another characteristic, conserved in both organisms, is protein folding, through the interaction of L-CK1.2 with chaperone proteins (Figure 5B). We have shown that L-CK1.2 interacts with *Leishmania* chaperone proteins, including HSP70s, HSP60, DNAJs, HSP110. Similarly in the macrophage, L-CK1.2 interacts with DNAJs, Hspa5 (Hsp70) and Hsp90. Interestingly, while *Leishmania* HSP90 is phosphorylated by L-CK1.2 (20), only host Hsp90 seems to interact with L-CK1.2. Finally, to determine whether L-CK1.2 targets specific pathways in the parasite, we compared the biological processes associated with L-CK1.2 in the parasite and the host cell by combining the L-CKAP*host* and substratome datasets and retained those specific to the parasite (this work and (19), Figure S5C, Table S9). The GO terms for several biological processes are specifically enriched in L-CK1.2 parasite interactome, including ‘precursor large subunit ribosomal RNA maturation’, ‘TOR signalling’ or ‘cell cycle regulation’ (Figure S5C, Table S9). Further investigations will be needed to determine whether this specificity might arise from interactions with proteins absent in the host or from differences in regulatory mechanisms.

Overall, although this experiment was conducted *in vitro* and may therefore capture interactions that do not occur under physiological conditions, it generated a comprehensive catalogue of potential host interacting partners of L-CK1.2. Notably, four proteins, Rcn2, Myo1f, Myo9b, and Hspa5, were identified in both L-CK1.2 host interactome and substratome, making them particularly attractive targets for further functional investigation (19).

## DISCUSSION

Currently, little is known about *Leishmania* CK1.2, despite its crucial role in parasite survival and its validation as a drug target (13, 18, 31), which might be linked to the challenges associated with studying an essential signalling kinase. Our findings shed light on the versatile and essential nature of L-CK1.2, demonstrated by its diverse subcellular localization and broad spectrum of interacting partners. We showed that the known localisation of L-CK1.2 interacting partners is consistent with L-CK1.2 distribution, suggesting that these proteins may confer the necessary specificity for its functions. This spatial coordination is particularly important given that L-CK1.2 is constitutively active (18). We identified potential roles for L-CK1.2 in the parasite, including its involvement in translational regulation, as evidenced by its interactions with numerous ribosomes and its involvement in molecular transport across the nuclear envelope. Additionally, specifically in axenic amastigotes, L-CK1.2 might play a role in vesicular trafficking, particularly endocytosis through its interaction with AP2 adaptor protein complex. We are currently investigating this hypothesis.

We also identified host biological processes that may be regulated by L-CK1.2 through interactions with host proteins, providing first insights into its potential role during infection. Many of the enriched host processes in our dataset, including cell death, immune response, and transport, are known to be modulated during *Leishmania* infection (69–72). The two main host processes targeted by L-CK1.2 seem to be ‘purine biosynthesis’’, more specifically linked to ‘glycolytic process’, and mitochondrial electron transport’, indicating the potential involvement of L-CK1.2 in the regulation of the host respiratory metabolism. This finding is consistent with recent data from Zhang *et al.* showing that macrophages infected with *Leishmania amazonensis* shift towards aerobic glycolysis for ATP production (67). Second, we identified many proteins linked to mitochondrial electron transport, particularly from the complex I, such as Ndufa10, Ndufs1 and Ndufs3, subunits of the mitochondrial membrane respiratory chain NADH dehydrogenase, important to maintain the mitochondrial membrane potential. *Leishmania* is known to target host mitochondria to modulate survival and also to control mitochondrial bioenergetics (73). Consistent with a role for L-CK1.2 in this process during *Leishmania* infection, human CK1δ/CK1ε have been involved in the regulation of complexes I and IV of the electron transport chain (74). The release of L-CK1.2 in the host cell via extracellular vesicles, as well as its ability to interact with and phosphorylate host proteins, suggest a dual role for L-CK1.2, in both the parasite and the host cell to regulate intracellular parasite survival (this work, (15, 19, 22)). Unlike viruses or bacteria that hijack host CK1 to manipulate cellular functions, *Leishmania* secretes its own CK1, suggesting that L-CK1.2 can functionally replace its mammalian counterparts. Supporting this hypothesis, our previous work demonstrated that the first 325 of the 353 amino acids, encompassing the catalytic domain and part of the C-terminus, are fully conserved among human-infecting *Leishmania* species, establishing L-CK1.2 as the most conserved kinase across these parasites (18). This conservation, unaffected by geographical diversity or differences in insect vectors, is likely driven by strong selective pressure from mammalian host cells on the L-CK1.2 protein sequence, explaining its high similarity to human orthologs (12, 18). However, this similarity does not extend to the C-terminus, suggesting that this domain might not be important for L-CK1.2 host functions. This hypothesis is not supported by the high conservation of L-CK1.2 C-terminal domain in most *Leishmania* species infecting humans (12, 18). One possible explanation lies in the presence of Low Complexity Regions (LCRs) within this domain. LCRs have been shown to be more abundant in highly connected proteins, such as signaling kinases, and contribute to protein interactions (75). Notably, LCRs are also found in the C-terminal domains of human CK1α, CK1δ, CK1ε, CK1γ1, CK1γ2, and CK1γ3 (Figure S6, http://smart.emblheidelberg.de/smart/set_mode.cgi?NORMAL=1, (76)). This conservation is striking, given the divergence of the L-CK1.2 C-terminus from its mammalian counterparts, suggesting that LCRs may be the key elements enabling L-CK1.2 to recognize proteins in the parasite and in the host. These protein-protein interactions are crucial, as L-CK1.2 is devoid of signal peptide or any similar domains (12, 18), and thus likely relies on host interacting partners to reach the subcellular localisation to exert its functions, like its mammalian orthologs(17). Conceivably, blocking these interactions might prevent L-CK1.2 functions in the host cell likely reducing parasite survival, a possibility we are currently investigating.

Another puzzling question is why *Leishmania* parasites would release an ortholog of kinases already present in the host cell, rather than subvert host CK1 like viruses or bacteria (77, 78)? Is L-CK1.2 in competition with the host CK1s? We hypothesize that during *Leishmania* infection, the functions targeted by L-CK1.2 might be attenuated by the inhibition or the reduced abundance of the host CK1s. One publication supports this hypothesis: Isnard *et al.* showed that CK1α, presents in the nucleus of host macrophage, decreases in abundance during *L. major* or *L. mexicana* infection (79). Interestingly, human CK1α phosphorylates IFNAR1/2 leading to its degradation (19, 22) and the removal of the interferon α and β receptor from the plasma membrane. This example shows that during infection while the macrophage would need to decrease CK1α to maintain those receptors at the cell surface and perceive the extracellular environment, the parasite would rather remove them by secreting its own CK1 to ensure that the macrophage is unaware of its environment, thus creating a safe niche for the parasite. Moreover, whereas the mammalian Ck1α, Ck1δ, and Ck1ε are autoinhibited and need the interactions with other proteins, such as Ddx3, for its activation, L-CK1.2 does not require interacting partners for its activity (18, 80).

One conserved feature of L-CK1.2 between the parasite and the host is its interaction with heat shock proteins, suggesting that L-CK1.2 might particularly contribute to the response to environmental cues. We, as well as other previously published studies, clearly show a strong link between L-CK1.2 and heat shock proteins of the parasite. L-CK1.2 is in complex with the *Leishmania* HSP70s and phosphorylates HSP23, P23 and HSP90, (this study, (20, 21)). The regulation of HSP23 and p23 is linked to temperature tolerance, while the phosphorylation of Hsp90 on Ser289 is important for parasite growth, flagella length and morphology (20, 21). Unexpectedly, we did not identify HSP90 among the L-CKAPs despite its abundance or any proteins involved in the foldosome, such as STI1 (81). This finding contrasts with human CK1s, which phosphorylate Hsp70 and Hsp90 to regulate the balance between protein folding and degradation (82), or whose abundance is regulated by Hsp90 (83). Noticeably, while *Leishmania* HSP90 was not identified among the L-CKAPs, host Hsp90ab1 was identified among the L-CKAP*host* (this work and (19)), suggesting that its HSP90-related functions may differ between the parasite and the macrophage, potentially resembling those of mammalian CK1s more closely. The importance of L-CK1.2 interaction with *Leishmania* HSP70 is unclear. HSP70 is part of a super family with several paralogs displaying different localisations such as the cytoplasm (HSPA1A), the lumen of the endoplasmic reticulum (HSPA5) or the mitochondrion (HSPA9A) (84). According to our data, L-CK1.2 interacts with several *Leishmania* HSP70 paralogs, although assigning a paralog was impossible due to the high percentage of identity between the different members. L-CK1.2 might also interact with host Hspa5, an endoplasmic reticulum chaperone involved in protein folding and quality control.

Our study provides extensive amount of data but has certain limitations, particularly regarding the host dataset, which was derived from macrophage lysates and does not account for localization constraints. Despite this limitation, protein class ontology analysis shows that our host dataset aligns well with the parasite dataset, which was obtained by tagging L-CK1.2 within the parasite. Future research will focus on validating these interactions in more physiological contexts, such as a cellular model or infected macrophages. Tracking the localization of L-CK1.2 in macrophages would be highly informative, though challenging due to its high similarity to host CK1s. We already know that the mouse anti-CK1 antibody cross-reacts with L-CK1.2, and our attempts to use a custom-made L-CK1.2 antibody for immunofluorescence have been unsuccessful. Additionally, tagging CK1.2 in the parasite with GFP or similarly sized fluorophores has proven ineffective, as it disrupts kinase function, limiting our options.

## CONCLUSION

Studying signalling kinases is inherently challenging, as their impairment affects multiple pathways, making it difficult to dissect precise regulatory mechanisms. Our study addresses this challenge by identifying specific interacting partners of L-CK1.2, which will be instrumental in elucidating its diverse functions in the parasite. Our findings highlight CK1.2 as a pivotal enzyme required not only for the survival of *Leishmania* but also for its successful adaptation to the host cell environment. We propose that this parasite-encoded serine/threonine kinase exerts dual functionality, acting in cis within the parasite itself while simultaneously operating in trans within the macrophage. In doing so, CK1.2 might establish itself as a bidirectional signalling hub that coordinates communication between pathogen and host. These findings position CK1.2 as a critical molecular effector that drives macrophage subversion and ultimately facilitates the establishment and persistence of infection ((15), this work). Furthermore, given the close relationship between *Leishmania* CK1.2 and its orthologs in *Trypanosoma brucei* (TbCK1.2), *Trypanosoma cruzi* (TcCK1.2), *Toxoplasma gondii* (TgCK1α), and *Plasmodium falciparum* (PfCK1)(18), our findings are likely to have broader implications for other parasitic infections.

## Acknowledgments

The authors would like to thanks Hira L Nakhasi, U.S. FDA, for the anti-*Ld*centrin antibody; Philippe Bastin, Institut Pasteur, for the anti-*Tb*IFT172 and anti-*Tb*PFR2 (L8C4) antibodies; Keith Gull, University of Oxford for the L1C6 antibody; the Unit of Technology and Service - Photonic BioImaging (UTechS PBI) from the Institut Pasteur and the Image Analysis Hub of the Institut Pasteur for the help with confocal microscopy and analyses of co-localisations, in particular Audrey Salles, Julien Fernandes, Anne Danckaert and Jean-Yves Tinevez. Finally, we would like to thank Brice Rotureau, Thierry Blisnick and Philippe Bastin for fruitful discussions and advice.

## Author contribution

Conceptualization: NR; Investigation: DM, OL and NR; Formal analysis: DM, OL, FD, VL, DL and NR; Writing - original draft: DM and NR; Writing - review & Editing: DM, OL, FD, VL, DL, GFS and NR; Funding acquisition: GFS and NR; Supervision NR.

