## Supplemental to Figures 1-5 for "Characterization of the dual functions of *Leishmania* CK1.2 in both the parasite and the macrophage"

### Figure S1

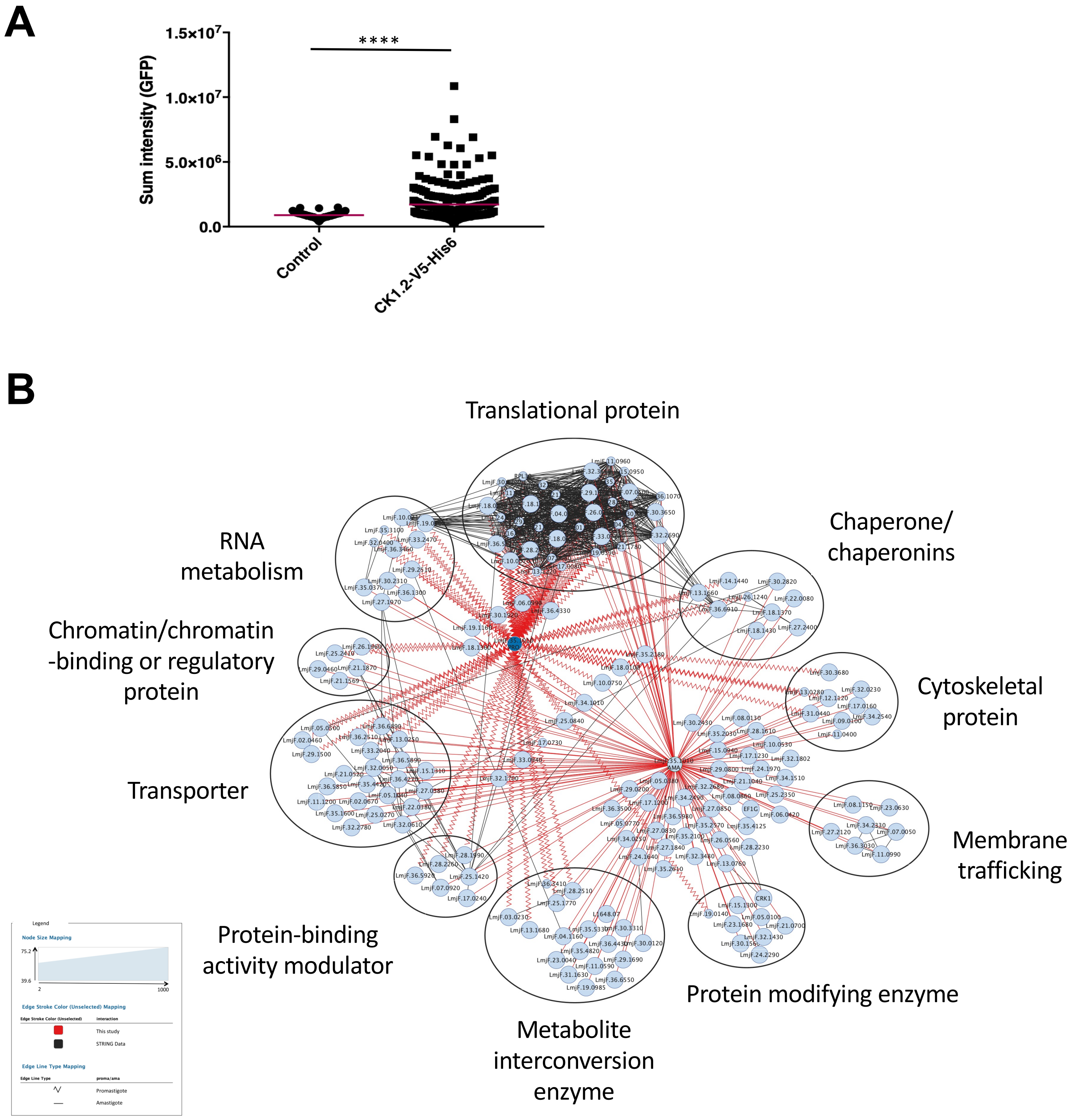


**Figure S1: L-CK1.2 has a pleiotropic localisation in the parasite**

**(A)** Analysis of different parameters extracted from ROI of the promastigotes parasite bodies. Scatter dot plots showing the sum of fluorescence intensity (for the V5 signal) of CK1.2-V5-expressing or mock control cell lines. The red line corresponds to the mean intensity. **(B)** Graphical representation of the affinity purification merged with the string app to visualise protein-protein interactions for PRO (bleu node and zigzag lines) and AxAMA (light blue node and straight line) using cytoscape app. The red lines represent protein-protein interaction identified in this study, while the black lines represent the protein-protein interactions identified in the STRING app. The size of the node represents the fold change ratio CK1.2/mock from 2 to 1000 (representing the peptides found in one condition.

### Figure S2

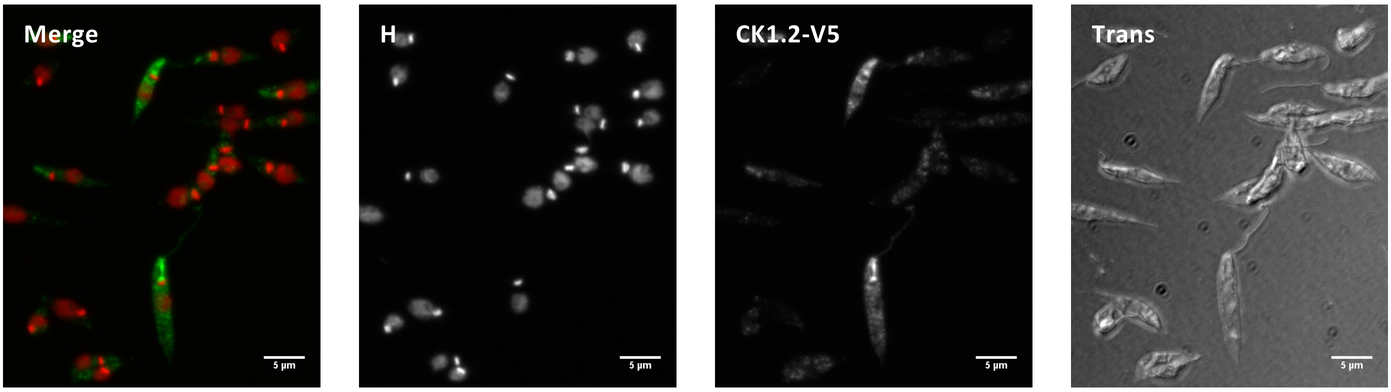


**Figure S2: CK1.2-V5 localisation in methanol-fixed promastigotes.**
IFA of *Ld*Bob pLEXSY-CK1.2-V5 **(A)** and *Ld*Bob pLEXSY **(mock, B)** promastigotes, fixed in ice-cold methanol for 3 minutes and stained with anti-V5 antibody to detect CK1.2-V5 localisation. The epifluorescence images were acquired under the same conditions and show the anti-V5 staining (CK1.2-V5 or V5), Hoechst 33342 staining (H), a merge of the anti-V5 (green) and H (red) signals and the transmission image (Trans). Scale bar, 5 µm. The pictures are maximum intensity projection of the z-stacks containing the parasites. The images presented in Figure S1 are representatives of at least 3 replicates.

### Figure S3

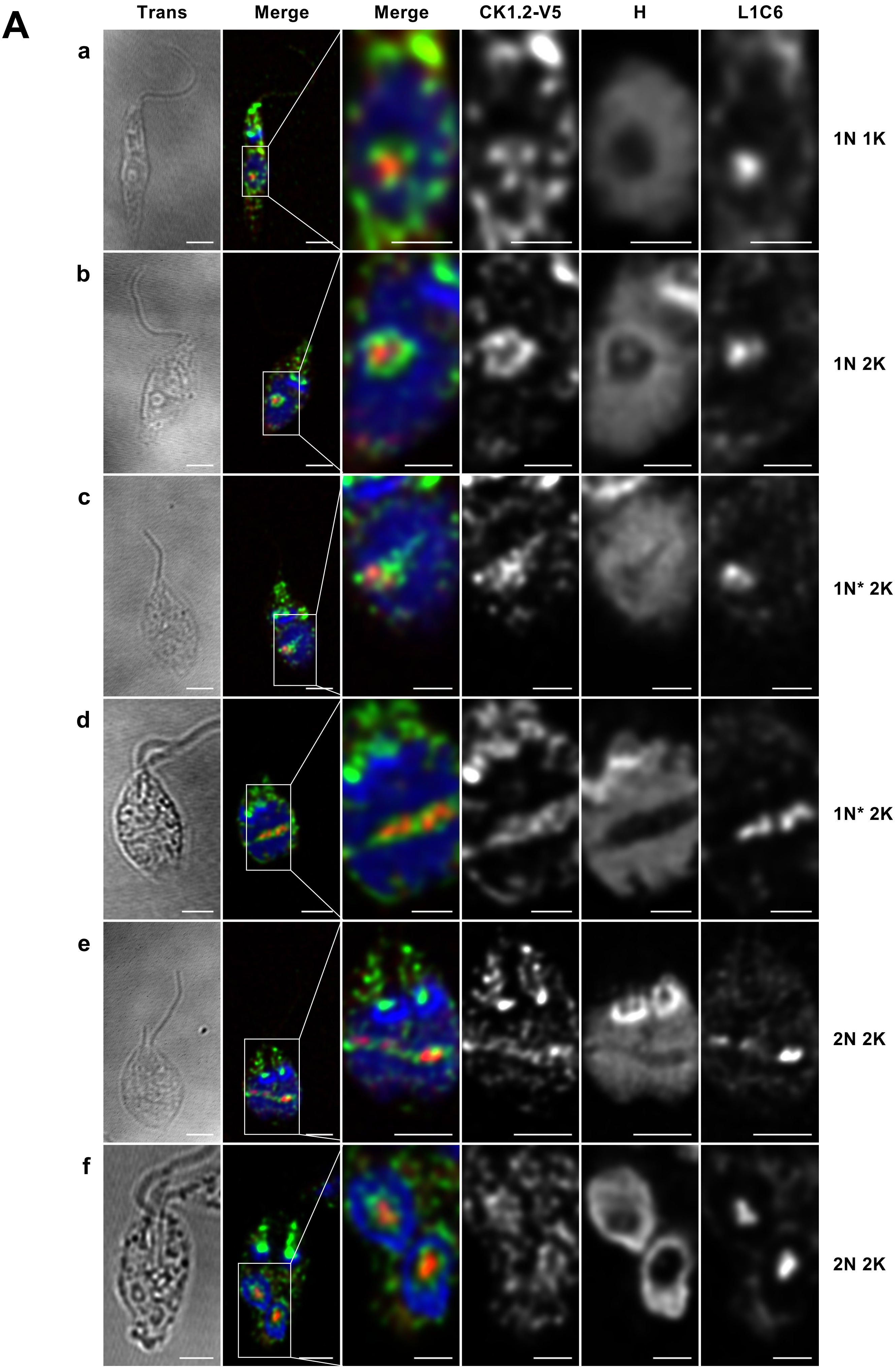
A
Trans
Merge
Merge
CK1.2-V5
H
L1C6
a
b
c
d
e
f
1N 1K
1N 2K
1N* 2K
1N* 2K
2N 2K
2N 2K

### Figure S3

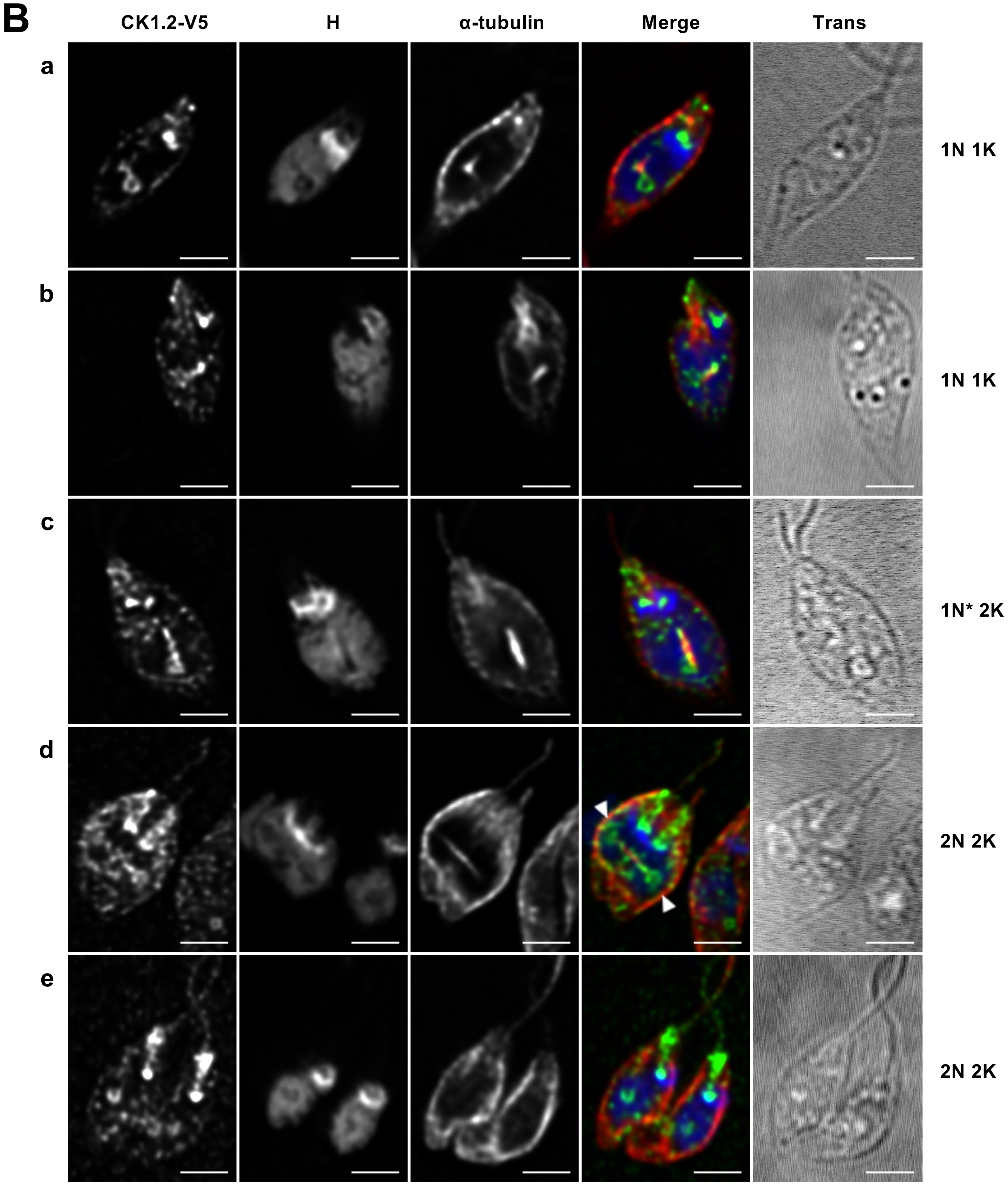
B
CK1.2-V5
H
α-tubulin
Merge
Trans
a
b
c
d
e
1N 1K
1N 1K
1N* 2K
2N 2K
2N 2K

**Figure S3: CK1.2 localises to the nucleolus and the mitotic spindle in the parasite.**

**(A)** IFA pictures of *Ld*Bob pLEXSY-CK1.2-V5 promastigotes obtained after detergent treatment followed by PFA fixation and stained with anti-V5 (CK1.2-V5) and anti-L1C6 (nucleolus, L1C6) antibodies. Confocal images representing sequential events of mitosis revealed different localisation patterns of L1C6 nucleolar marker and CK1.2-V5. (a – f) The images correspond to the transmission (Trans), the merged containing CK1.2-V5 (green), Hoechst 33342 (H) (blue) and L1C6 (red) signals. The following four images show a magnification of the nuclear region with the merged and single channel images. N=nucleus, K=kinetoplast. Scale bar, 2 µm or 1 µm for magnified images. These pictures are single stacks extracted from deconvolved confocal stacks corrected for chromatic aberration. **(B)** IFA pictures of *Ld*Bob pLEXSY-CK1.2-V5 promastigotes obtained after detergent treatment followed by PFA fixation and stained with anti-V5 and anti-α-tubulin antibodies. Sequential images of various stages of cell division (a – e) showing the single channel images for CK1.2-V5, H and α-tubulin signals, the merged images showing CK1.2-V5 (green), H (blue) and α-tubulin (red) signals, and the transmission image (Trans). Scale bar, 2 µm. These pictures are single stacks extracted from deconvolved confocal stacks corrected for chromatic aberration. The images presented in Figure S3 are representatives of at least 3 replicates. **(C)** Schematic of the nuclear pore complex adapted from Obado *et al.* {Obado, 2016 #526} and the proteins identified in L-CK1.2 immuno-precipitation (Figure 1B). Red border: proteins identified in axenic amastigotes; Blue border: proteins identified in promastigotes; and grey square with red border: protein added compared to the schematic published by Obado *et al.* {Obado, 2016 #526}.

### Figure S4

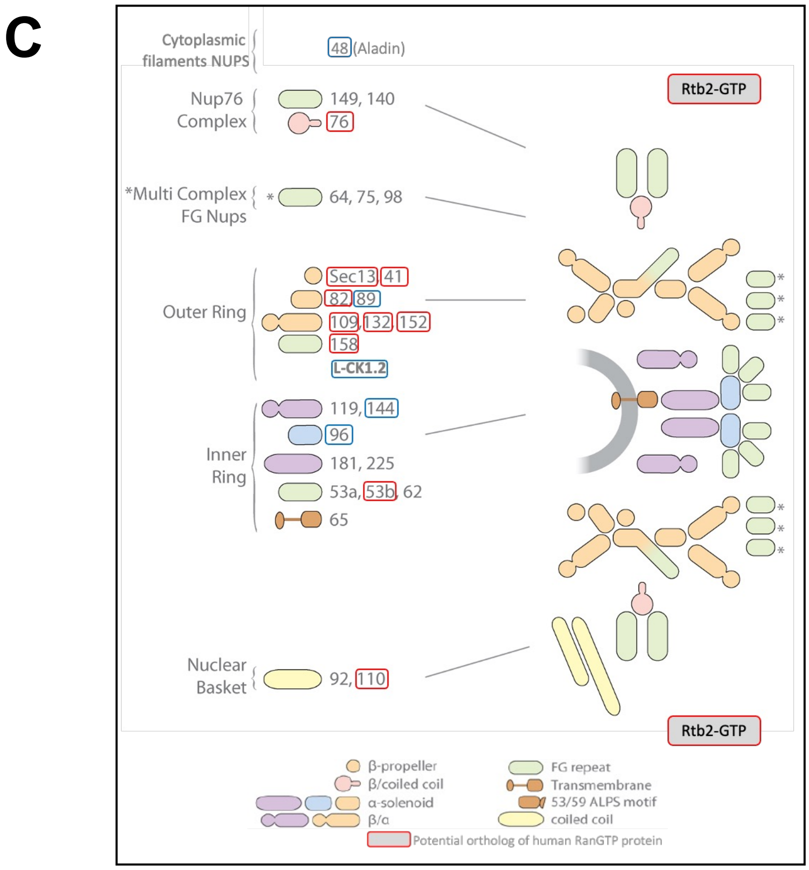


**Figure S4: CK1.2 interacts with the nuclear pore complexes.**
Schematic adapted from Obado *et al.* {Obado, 2016 #526}. Red border: proteins identified in axenic amastigotes; Blue border: proteins identified in promastigotes; and grey square with red border: protein added compared to the schematic published by Obado *et al.* {Obado, 2016 #526}.

### Figure S5

**A**

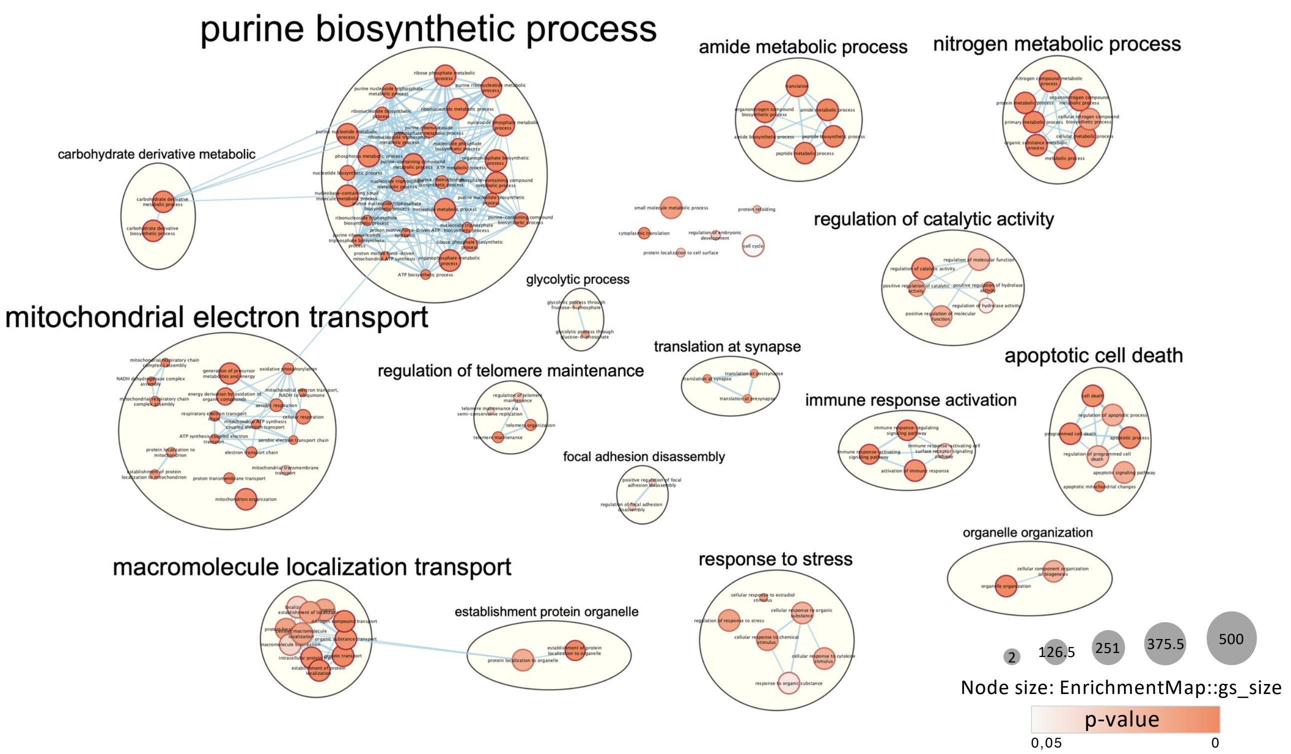
purine biosynthetic process
amide metabolic process
nitrogen metabolic process
carbohydrate derivative metabolic
regulation of catalytic activity
glycolytic process
mitochondrial electron transport
translation at synapse
regulation of telomere maintenance
apoptotic cell death
immune response activation
focal adhesion disassembly
organelle organization
macromolecule localization transport
response to stress
establishment protein organelle
2
126.5
251
375.5
500
Node size: EnrichmentMap::gs_size
p-value
0,05
0

**B**

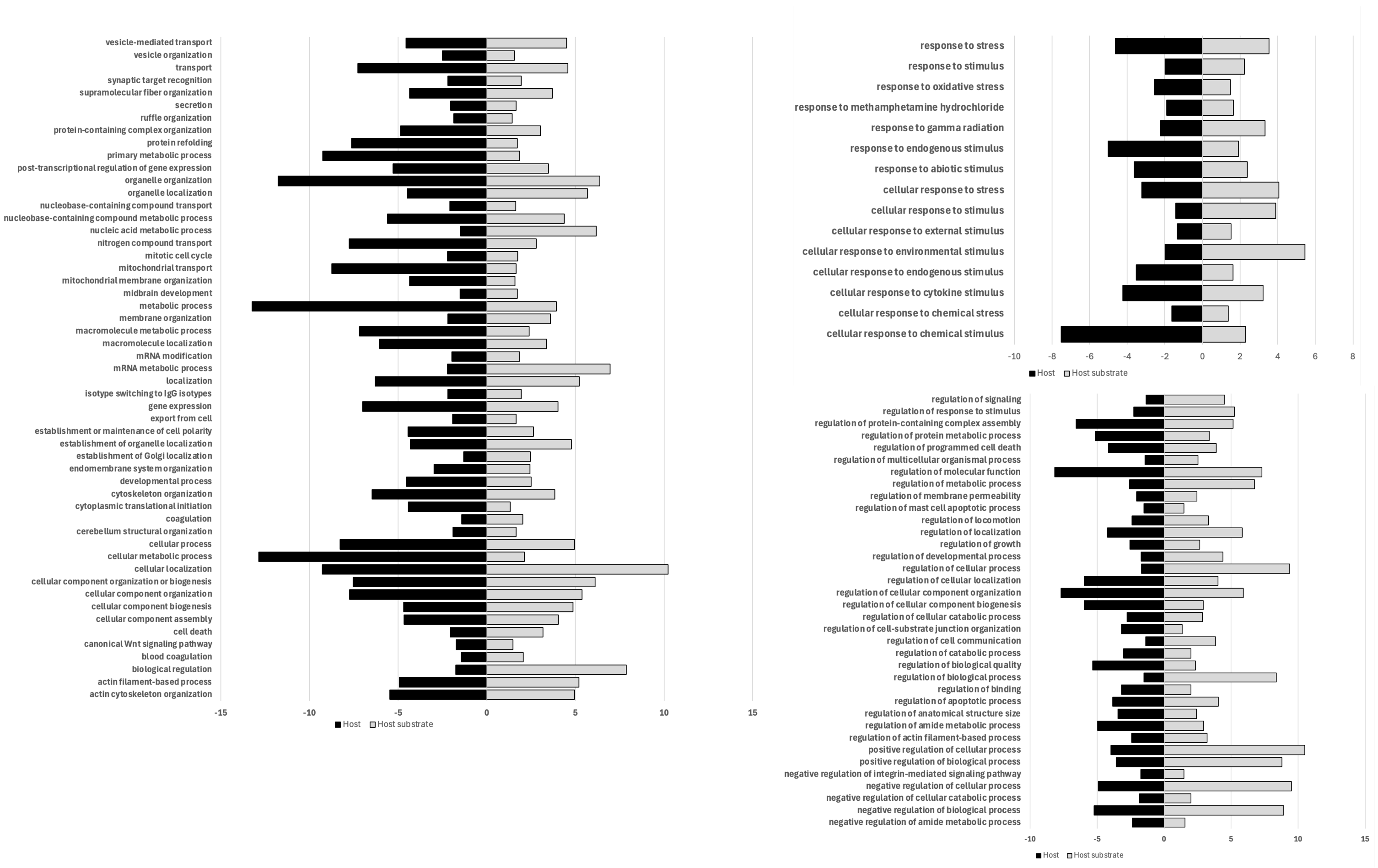
vesicle-mediated transport
vesicle organization
transport
synaptic target recognition
supramolecular fiber organization
secretion
ruffle organization
protein-containing complex organization
protein refolding
primary metabolic process
post-transcriptional regulation of gene expression
organelle organization
organelle localization
nucleobase-containing compound transport
nucleobase-containing compound metabolic process
nucleic acid metabolic process
nitrogen compound transport
mitotic cell cycle
mitochondrial transport
mitochondrial membrane organization
midbrain development
metabolic process
membrane organization
macromolecule metabolic process
macromolecule localization
mRNA modification
mRNA metabolic process
localization
isotype switching to IgG isotypes
gene expression
export from cell
establishment or maintenance of cell polarity
establishment of organelle localization
establishment of Golgi localization
endomembrane system organization
developmental process
cytoskeleton organization
cytoplasmic translational initiation
coagulation
cerebellum structural organization
cellular process
cellular metabolic process
cellular localization
cellular component organization or biogenesis
cellular component organization
cellular component biogenesis
cellular component assembly
cell death
canonical Wnt signaling pathway
blood coagulation
biological regulation
actin filament-based process
actin cytoskeleton organization
-15
-10
-5
0
5
10
15
Host
Host substrate
response to stress
response to stimulus
response to oxidative stress
response to methamphetamine hydrochloride
response to gamma radiation
response to endogenous stimulus
response to abiotic stimulus
cellular response to stress
cellular response to stimulus
cellular response to external stimulus
cellular response to environmental stimulus
cellular response to endogenous stimulus
cellular response to cytokine stimulus
cellular response to chemical stress
cellular response to chemical stimulus
-10
-8
-6
-4
-2
0
2
4
6
8
Host
Host substrate
regulation of signaling
regulation of response to stimulus
regulation of protein-containing complex assembly
regulation of protein metabolic process
regulation of programmed cell death
regulation of multicellular organismal process
regulation of molecular function
regulation of metabolic process
regulation of membrane permeability
regulation of mast cell apoptotic process
regulation of locomotion
regulation of localization
regulation of growth
regulation of developmental process
regulation of cellular process
regulation of cellular localization
regulation of cellular component organization
regulation of cellular component biogenesis
regulation of cellular catabolic process
regulation of cell-substrate junction organization
regulation of cell communication
regulation of catabolic process
regulation of biological quality
regulation of biological process
regulation of binding
regulation of apoptotic process
regulation of anatomical structure size
regulation of amide metabolic process
regulation of actin filament-based process
positive regulation of cellular process
positive regulation of biological process
negative regulation of integrin-mediated signaling pathway
negative regulation of cellular process
negative regulation of cellular catabolic process
negative regulation of biological process
negative regulation of amide metabolic process
-10
-5
0
5
10
15
Host
Host substrate

C

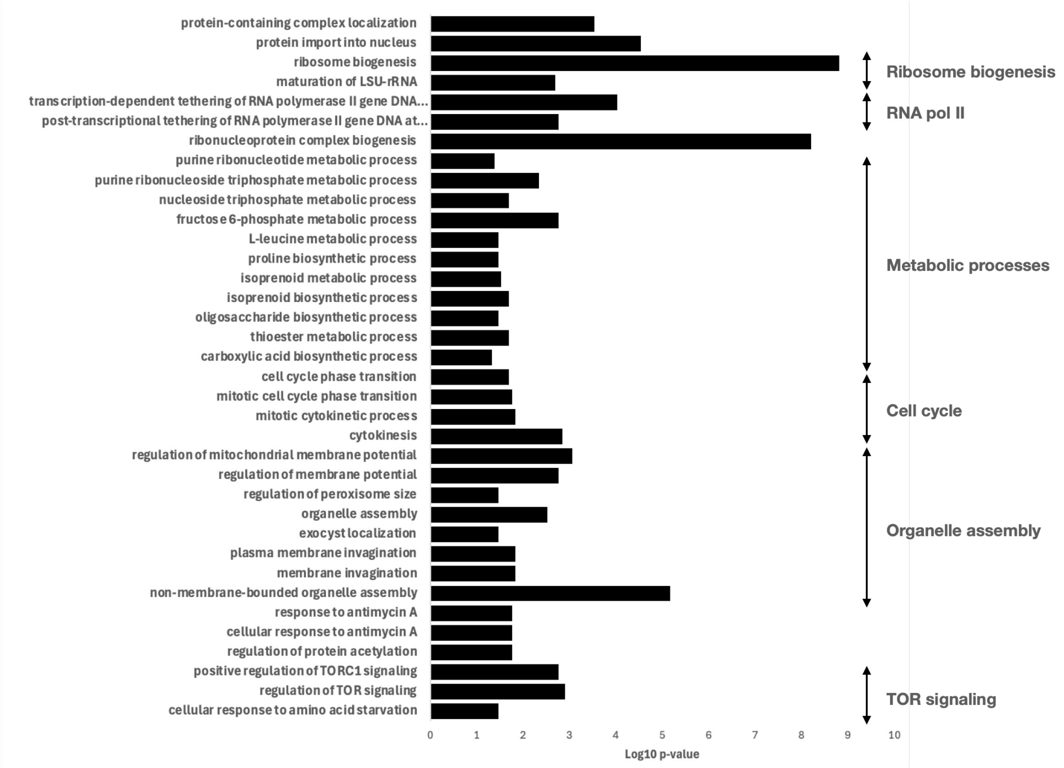


**Figure S5: L-CK1.2 interacts with macrophage proteins.**
**(A)** Biological process GO term enrichment in the L-CKAP*Host* dataset. The scale font of the clusters represents their size, the node colour represents the p-value, and the size of the node represents the enrichmentmap gs_size. **(B)** GO term enrichment for *Leishmania* specific host 'biological process' common between the interactome (this study) and the substratome {Smirlis, 2021 #470} of L-CK1.2. It was performed using Tritrypdb-Revigo and the cytoscape app. **(C)** Specific GO term enrichment of in the interactome of L-CK1.2 in the parasite. Difference in GO term enrichment for biological processes between L-CKAP dataset and combined L-CKAP*Host*+L-CK1.2 host substrates dataset.

### Figure S6

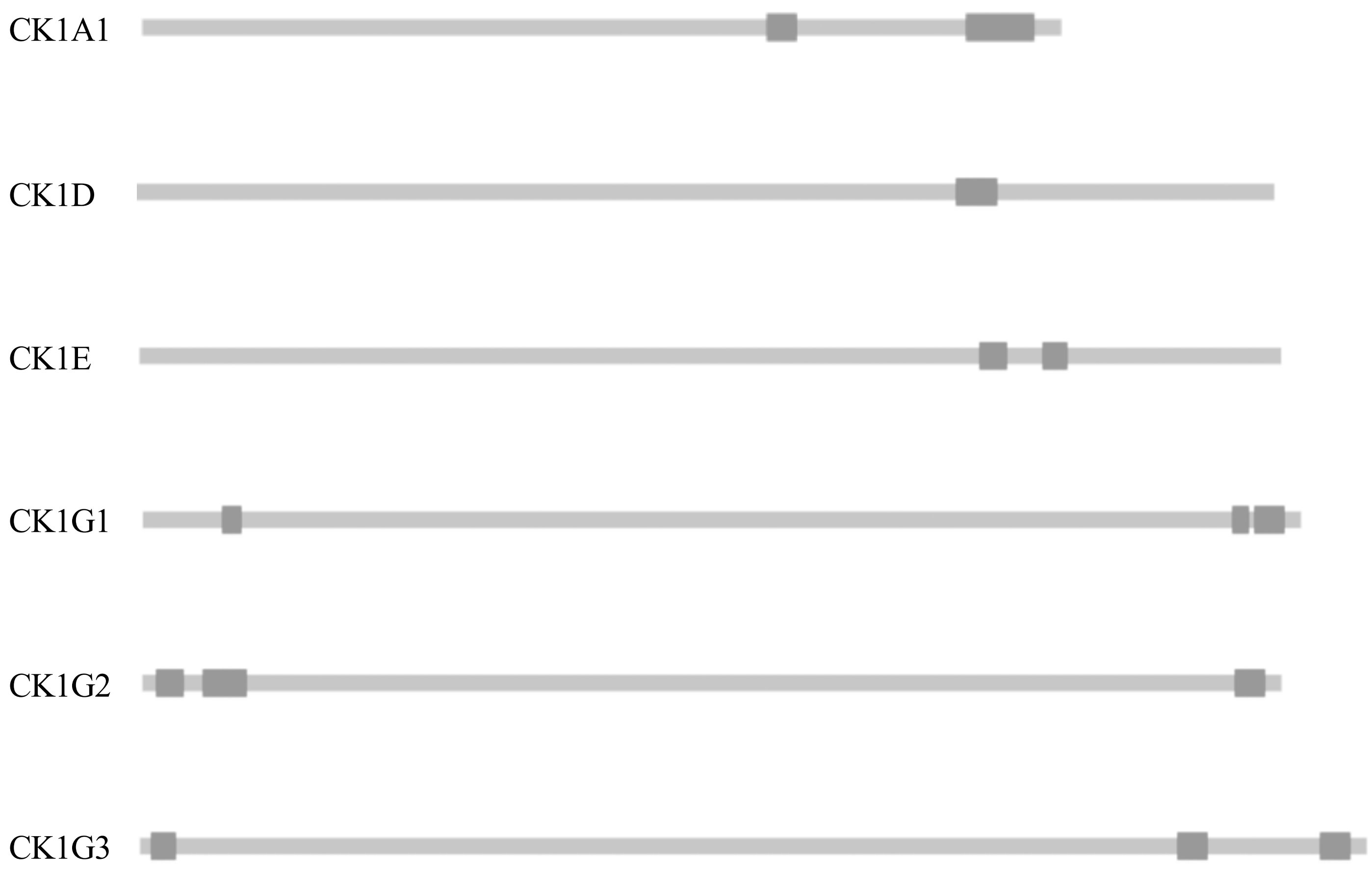


**Figure S6: Position of the LCR in human CK1s.**

Cartoon representing the low complexity region (LCR, dark grey) on the protein sequence (light grey) of human CK1. CK1A (P48729), CK1D (P48730), CK1E (P49674), CK1G1 (Q9HCP0), CK1G2 (P78368) and CK1G3 (Q9Y6M4).

Table S3: protein class

| 1st level | Gene number | |
|---|---|---|
| | PRO | AxAMA |
| protein modifying enzyme (PC00260) | 2 | 10 |
| transporter (PC00227) | 5 | 11 |
| membrane traffic protein (PC00150) | 0 | 7 |
| chaperone (PC00072) | 6 | 7 |
| DNA metabolism protein (PC00009) | 0 | 1 |
| protein-binding activity modulator (PC00095) | 1 | 4 |
| RNA metabolism protein (PC00171) | 4 | 4 |
| cytoskeletal protein (PC00085) | 1 | 5 |
| gene-specific transcriptional regulator (PC00264) | 1 | 1 |
| translational protein (PC00263) | 33 | 17 |
| metabolite interconversion enzyme (PC00262) | 5 | 14 |
| chromatin-binding, or -regulatory protein (PC00077) | 1 | 4 |

Table S4: Biological processes

| 1 st level | 2nd level | Gene nb | |
|---|---|---|---|
| | | PRO | AxAMA |
| biological regulation (GO:0065007) | regulation of biological process (GO:0050789) | 4 | 8 |
| | regulation of biological quality (GO:0065008) | 2 | 2 |
| cellular process (GO:0009987) | cellular component organization or biogenesis (GO:0071840) | 16 | 23 |
| | cell communication (GO:0007154) | 1 | 3 |
| | cellular metabolic process (GO:0044237) | 18 | 23 |
| | Microtubule-based process (GO:0007017) | 1 | 3 |
| | Signal transduction (GO:0007165) | 1 | 3 |
| | export from cell (GO:0140352) | 0 | 2 |
| | exocytic process (GO:0140029) | 0 | 2 |
| | cell cycle process (GO:0022402) | 1 | 2 |
| | cellular homeostasis (GO:0019725) | 0 | 1 |
| | cellular localization (GO:0051641) | 5 | 11 |
| | cellular response to stimulus (GO:0051716) | 2 | 5 |
| | transmembrane transport (GO:0055085) | 1 | 2 |
| | protein folding (GO:0006457) | 3 | 5 |
| | vesicle-mediated transport (GO:0016192) | 0 | 8 |
| | cell cycle (GO:0007049) | 1 | 2 |
| | movement of cell or subcellular component (GO:0006928) | 0 | 1 |
| | process utilizing autophagic mechanism (GO:0061919) | 0 | 1 |
| localization (GO:0051179) | macromolecule localization (GO:0033036) | 4 | 7 |
| | establishment of localization (GO:0051234) | 6 | 16 |
| | cellular localization (GO:0051641) | 5 | 11 |
| | protein-containing complex localization (GO:0031503) | 0 | 1 |
| metabolic process (GO:0008152) | small molecule metabolic process (GO:0044281) | 3 | 5 |
| | biosynthetic process (GO:0009058) | 13 | 10 |
| | ATP metabolic process (GO:0046034) | 1 | 0 |
| | nitrogen compound metabolic process (GO:0006807) | 17 | 19 |
| | catabolic process (GO:0009056) | 0 | 3 |
| | primary metabolic process (GO:0044238) | 18 | 21 |
| | cellular metabolic process (GO:0044237) | 18 | 23 |
| | organic substance metabolic process (GO:0071704) | 19 | 22 |
| response to stimulus (GO:0050896) | response to external stimulus (GO:0009605) | 0 | 1 |
| | response to stress (GO:0006950) | 1 | 3 |
| | response to chemical (GO:0042221) | 1 | 1 |
| | cellular response to stimulus (GO:0051716) | 2 | 5 |

Table S5

| Localisation | Corresponding genes |
| --- | --- |
| **Protein/vesicular trafficking** | 60S ribosomal protein L21 (LdBPK_343440.1); Nodulin-like/ Pyruvate transporter (LdBPK_291600.1); LdBPK_071070.1; Organic solute transporter Ostalpha (LdBPK_111190.1); C2 domain containing protein (LdBPK_130650.1); LdBPK_171330.1; heat shock protein DNAJ (LdBPK_272350.1); cyclin dependent kinase-binding protein (LdBPK_282390.1); Gem-associated protein 2 (LdBPK_361350.1). |
| **Nucleolus** | 40S ribosomal protein S7 (LdBPK_010430.1), S5 (LdBPK_110960.1), S4 (LdBPK_131120.1), S6 (LmjF.21.1780), S11 (LdBPK_211790.1), S13 (LdBPK_333300.1), S9 (LdBPK_361310.1) ; 60S ribosomal protein L11 (LdBPK_220004.1), L21 (LdBPK_343440.1), L7 (LdBPK_180230.1/LdBPK_260160.1), L10a (LdBPK_363950.1), L34 (LdBPK_363930.1), L26 (LdBPK_242140.1), L1a (LdBPK_291160.1), L13 (LdBPK_292570.1,LdBPK_292580.1), L18a (LdBPK_350600.1), L6 (LdBPK_330770.1), L30 (LdBPK_350240.1), L7a (LdBPK_070550.1), L27 (LdBPK_322830.1); nucleolar protein 56 (LdBPK_100210.1); alpha tubulin (LdBPK_130330.1); fibrillarin (LdBPK_190090.1,LdBPK_363220.1); LdBPK_252520.1; T-complex protein 1, theta subunit (LdBPK_367240.1); histone deacetylase (LdBPK_081300.1); Nucleoporin NUP41 (LdBPK_210580.1); heat shock protein DNAJ (LdBPK_220009.1); class I transcription factor A, subunit 1 (LdBPK_270681.1). |
| **Nucleus / kinetoplast / antipodal sites** | nucleolar protein 56 (LdBPK_100210.1); LdBPK_180100.1; fibrillarin (LdBPK_190090.1, LdBPK_363220.1); 40S ribosomal protein S3A(LdBPK_350400.1), S9 (LdBPK_361310.1); 60S ribosomal protein L26 (LdBPK_242140.1); heat shock protein 70, putative,heat-shock protein hsp70, putative,heat shock protein 70-related protein,luminal binding protein 1 (LdBPK_261220.1,LdBPK_281310.1,LdBPK_283000.1,LdBPK_283040.1); Nodulin-like, putative / Pyruvate transporter (LdBPK_291600.1); LdBPK_341080.1; casein kinase I (LdBPK_351030.1); kinetoplast DNA-associated protein (LdBPK_366180.1); ATP synthase F1, alpha subunit (LdBPK_050510.1); Microtubule associated protein (MAP65/ASE1 family, LdBPK_151000.1); histone deacetylase (LdBPK_081300.1); Nucleoporin NUP41 (LdBPK_210580.1); heat shock protein DNAJ (LdBPK_220009.1); class I transcription factor A, subunit 1 (LdBPK_270700.1); Small nuclear ribonucleoprotein-associated protein B (snRNP-B, LdBPK_271890.1); MSP domain containing protein (LdBPK_272040.1); heat shock protein DNAJ (LdBPK_272350.1); SNF2 family N-terminal domain/Helicase conserved C-terminal domain containing protein (LdBPK_290210.1); dynein light chain (LdBPK_320240.1); nucleoside diphosphate kinase b (LdBPK_323110.1); LdBPK_363660.1; Mitoribosomal LSU assembly factor 24 (LdBPK_302320.1); mitochondrial phosphate transporter (LdBPK_354490.1); mitochondrial carrier protein (LdBPK_020640.1). |
| **Nuclear pore** | Nucleoporin NUP48 (LdBPK_130250.1); NUP89 (LdBPK_332160.1); NUP96 (LdBPK_362640.1); NUP144 (NUP155-like, LdBPK_367220.1); LdBPK_091010.1; LdBPK_100580.1; NUP41 (LdBPK_210580.1); NUP132 (LdBPK_220260.1); NUP76 (LdBPK_242050.1); NUP110 (LdBPK_250270.1); GTP-binding nuclear protein rtb2 (LdBPK_251460.1); NUP158 (LdBPK_270390.1); NUP53b (LdBPK_290830.1); NUP109 (LdBPK_322920.1); NUP82 (LdBPK_351600.1); LdBPK_281740.1; protein transport protein Sec13 (LdBPK_320050.1); NUP152 (LdBPK_364480.1). |
| **Flagella connector/hook complex/FAZ/microtubule quartet/flagellar pocket collar** | LdBPK_061020.1; LdBPK_181340.1; LdBPK_282120.1; FAZ 2 (LdBPK_120720.1); Associated kinase of Tb14-3-3 (LdBPK_190130.1); predicted C2 domain protein (LdBPK_364540.1); phosphoprotein phosphatase (KPP1, LdBPK_050100.1); Acyl-coenzyme A thioesterase (LdBPK_291820.1); dynein light chain, flagellar outer arm (LdBPK_320240.1); dual specificity protein phosphatase (LdBPK_210760.1); LdBPK_322820.1; Coiled-coil and C2 domain-containing protein (LdBPK_342370.1); MORN repeat-containing protein 1 (LdBPK_303360.1); LdBPK_352610.1; SUMO-interacting motif-containing protein (LdBPK_321490.1); bilbo1 (LdBPK_090120.1). |
| **Cortical cytoskeleton/Cell tip/membrane/pellicular membrane** | LdBPK_061020.1; alpha tubulin (LdBPK_130330.1); cytoskeleton associated protein (LdBPK_141540.1); LdBPK_191150.1; LdBPK_303740.1; cytoskeleton-associated protein CAP5.5 (LdBPK_310450.1); long chain fatty Acyl CoA synthetase (LdBPK_030220.1); FAZ 2 (LdBPK_120720.1); LdBPK_252520.1; casein kinase I (CK1.2, LdBPK_351030.1); microtubule-associated protein (LdBPK_050380.1) ; Dual specificity phosphatase, catalytic domain containing protein (LdBPK_271740.1); kharon1 (LdBPK_366110.1); dynein light chain, flagellar outer arm (LdBPK_320240.1); LdBPK_352610.1. |
| **Glycosome/acidocalcisome/ER/Golgi/lipid droplet** | glycosomal membrane protein (LdBPK_282430.1); ATP-dependent 6-phosphofructokinase (LdBPK_292620.1); long chain fatty Acyl CoA synthetase (LdBPK_030220.1); 3-oxo-5-alpha-steroid 4-dehydrogenase (LdBPK_251850.1); heat shock protein 70 (LdBPK_261220.1, LdBPK_281310.1, LdBPK_283000.1, LdBPK_283040.1); Nodulin-like, putative / Pyruvate transporter (LdBPK_291600.1); fructose-1,6-bisphosphatase (LdBPK_041170.1); alkyldihydroxyacetonephosphate synthase (LdBPK_300120.1); isopentenyl-diphosphate delta-isomerase (LdBPK_355300.1); MGT1 magnesium transporter (LdBPK_151330.1); protein transport protein SEC13 (LdBPK_320050.1); Histidine phosphatase superfamily (LdBPK_354880.1). |
| **Mitochondrion** | Mitochondrial outer membrane protein porin (LdBPK_020430.1); ATP synthase F1, alpha subunit (LdBPK_050510.1); LdBPK_061020.1; POMP11 / Present in the outer mitochondrial membrane proteome 11 (LdBPK_332600.1); mitochondrial carrier protein (LdBPK_020640.1); stomatin-like protein (LdBPK_051040.1); cell division protein kinase 2 (LdBPK_211320.1); Mitoribosomal LSU assembly factor 24 (LdBPK_302320.1); mitochondrial phosphate transporter (LdBPK_354490.1). |
| **Spindle** | Nucleus and spindle associated protein 2 (MAP65/ASE1 family, LdBPK_151000.1). |

### Table S5

| Localisation | Corresponding genes |
|---|---|
| **Basal body, TAC and TF** | FAZ 2 (LdBPK_120720.1); Associated kinase of Tb14-3-3 (LdBPK_190130.1); casein kinase I (LdBPK_351030.1); microtubule-associated protein (LdBPK_050380.1); Regulator of chromosome condensation (RCC1) repeat (LdBPK_170290.1); dual specificity protein phosphatase (LdBPK_210760.1); Nucleoporin NUP41 (LdBPK_210580.1); peroxidoxin (LdBPK_230050.1); dynein light chain, flagellar outer arm (LdBPK_320240.1); LdBPK_323680.1; Tripartite Attachment Complex Protein 60 (LdBPK_260530.1); heat shock protein DNAJ (LdBPK_220009.1). |
| **Flagellar pocket** | long chain fatty Acyl CoA synthetase (LdBPK_030220.1); LdBPK_252520.1; Exocyst complex component Sec3 (LdBPK_081060.1); Exocyst complex component EXO99 (LdBPK_211910.1); exocyst complex component Sec10 (LdBPK_230800.1); MSP (Major sperm protein) domain containing protein (LdBPK_272040.1); LdBPK_323680.1; ; alpha-adaptin-like protein (LdBPK_070060.1); adaptin-related protein-like protein (LdBPK_110990.1); clathrin coat assembly protein AP17 (LdBPK_342100.1); clathrin coat assembly protein-like protein (LdBPK_363180.1). |
| **Flagellum/Flagellar tip/paraflagellar rod** | LdBPK_180100.1; Associated kinase of Tb14-3-3 (LdBPK_190130.1); 60S ribosomal protein L26 (LdBPK_242140.1); 40S ribosomal protein S9 (LdBPK_361310.1); Nucleoporin NUP96 (LdBPK_362640.1); 40S ribosomal protein SA (LdBPK_365353.1); kinetoplast DNA-associated protein (LdBPK_366180.1); heat shock protein 70 (LdBPK_261220.1, LdBPK_281310.1, LdBPK_283000.1, LdBPK_283040.1); alpha tubulin (LdBPK_130330.1); Nucleoporin NUP41 (LdBPK_210580.1); peroxidoxin (LdBPK_230050.1); MSP (Major sperm protein) domain containing protein (LdBPK_272040.1); heat shock protein DNAJ (LdBPK_272350.1); dynein light chain, flagellar outer arm (LdBPK_320240.1); nucleoside diphosphate kinase b (LdBPK_323110.1); Chaperonin HSP60 (LdBPK_302830.1,LdBPK_321940.1,LdBPK_362130.1,LdBPK_362140.1); Inositol hexakisphosphate (LdBPK_231720.1). |
| **Cytoplasm** | 40S ribosomal protein S3 (LdBPK_330960.1), S4 (LdBPK_131120.1), S5 (LdBPK_110960.1), S6 (LmjF.21.1780), S7 (LdBPK_010430.1), S9 (LdBPK_361310.1), S11 (LdBPK_211790.1), S13 (LdBPK_333300.1), S17 (LdBPK_282750.1), S20 (LdBPK_281100.1); SA (LdBPK_365353.1); 60S ribosomal protein L1a (LdBPK_291160.1), L6 (LdBPK_330770.1), L7 (LdBPK_260160.1), L7a (LdBPK_070550.1), L10 (LdBPK_040750.1), L10a (LdBPK_363950.1); L11 (LdBPK_220004.1), L18a (LdBPK_350600.1), L21 (LdBPK_343440.1), L26 (LdBPK_242140.1), L24 (LdBPK_361130.1); L27 (LdBPK_322830.1); L28 (LdBPK_111110.1), L30 (LdBPK_350240.1), L34 (LdBPK_363930.1); alpha tubulin (LdBPK_130330.1); chaperonin TCP20 (LdBPK_131400.1); pyrroline-5-carboxylate reductase (LdBPK_131420.1); elongation factor 1-alpha (LdBPK_170170.1); LdBPK_180100.1; Associated kinase of Tb14-3-3 (LdBPK_190130.1); 3-oxo-5-alpha-steroid 4-dehydrogenase (LdBPK_251850.1); heat shock protein 70 (LdBPK_261220.1,LdBPK_281310.1,LdBPK_283000.1,LdBPK_283040.1); LdBPK_261960.1; acyl-CoA dehydrogenase (LdBPK_282700.1); Nodulin-like/ Pyruvate transporter (LdBPK_291600.1); ATP-dependent RNA helicase HEL67 (LdBPK_320410.1); casein kinase I (LdBPK_351030.1); Nucleoporin NUP96 (LdBPK_362640.1); C2 domain protein (LdBPK_130650.1, LdBPK_364540.1); kinetoplast DNA-associated protein (LdBPK_366180.1); LdBPK_071070.1; Microtubule associated protein (MAP65/ASE1 family, LdBPK_151000.1); ubiquitin hydrolase (LdBPK_151320.1); MGT1 magnesium transporter (LdBPK_151330.1); Regulator of chromosome condensation (RCC1) repeat (LdBPK_170290.1); LdBPK_171330.1; heat shock protein 110 (LdBPK_181350.1); kinesin (LdBPK_211280.1); dual specificity protein phosphatase (LdBPK_210760.1); Nucleoporin NUP41 (LdBPK_210580.1); heat shock protein DNAJ (LdBPK_220009.1); peroxidoxin (LdBPK_230050.1); exocyst complex component Sec10 (LdBPK_230800.1); Male sterility protein (LdBPK_241710.1); Nucleoporin NUP76 (LdBPK_242050.1); Nucleoporin NUP110 (LdBPK_250270.1); GTP-binding nuclear protein rtb2 (LdBPK_251460.1); LdBPK_252520.1; Dual specificity phosphatase (LdBPK_271740.1); MSP (Major sperm protein) domain containing protein (LdBPK_272040.1); heat shock protein DNAJ (LdBPK_272350.1); LdBPK_281740.1; cyclin dependent kinase-binding protein (LdBPK_282390.1); Acyl-coenzyme A thioesterase (LdBPK_291820.1); alkyldihydroxyacetonephosphate synthase (LdBPK_300120.1); Mitoribosomal LSU assembly factor 24 (LdBPK_302320.1); dynein light chain, flagellar outer arm (LdBPK_320240.1); LdBPK_320640.1; LdBPK_322820.1; nucleoside diphosphate kinase b (LdBPK_323110.1); LdBPK_323680.1; ATP-dependent DEAD-box RNA helicase (LdBPK_350370.1); LdBPK_352610.1; Gem-associated protein 2 (LdBPK_361350.1); Chaperonin HSP60 (LdBPK_302830.1, LdBPK_321940.1, LdBPK_362130.1, LdBPK_362140.1); flagellum targeting protein kharon1 (LdBPK_366110.1). |
| **Unknown localisation** | LdBPK_100800.1; GRAM domain containing protein (LdBPK_170750.1); 40S ribosomal protein S23 (LdBPK_211300.1), S14 (LdBPK_303650.1), S26 (LdBPK_280570.1); LdBPK_250870.1; LdBPK_301900.1; LdBPK_321780.1; LdBPK_330990.1; 60S ribosomal protein L3 (LdBPK_323320.1), L15 (LdBPK_303710.1), L35a (LdBPK_342240.1); LdBPK_352220.1; ATP-dependent DEAD/H RNA helicase (LdBPK_353150.1); short chain dehydrogenase-like protein (LdBPK_363570.1); NIPSNAP (LdBPK_363620.1); LdBPK_050770.1; LdBPK_060440.1; LdBPK_080130.1; LdBPK_080800.1; LdBPK_100800.1; tubulin-tyrosine ligase-like protein(LdBPK_110400.1); 3-methylcrotonoyl-CoA carboxylase beta subunit (LdBPK_110600.1); kinesin motor domain containing protein (LdBPK_170270.1); LdBPK_171300.1; chaperone DNAJ protein (LdBPK_181410.1); 4-coumarate:coa ligase-like protein (LdBPK_190940.1); cullin-like protein-like protein (LdBPK_242380.1); LdBPK_252460.1; LdBPK_290470.1; protein kinase (LdBPK_301580.1); LdBPK_302440.1; 3-ketoacyl-CoA thiolase-like protein (LmjF.31.1630); LdBPK_321890.1; LdBPK_340270.1; LdBPK_341610.1; LdBPK_342300.1; LdBPK_352020.1; LdBPK_352100.1; LdBPK_352650.1 LdBPK_354190.1; asparaginase-like protein (LdBPK_364650.1); LdBPK_366150.1; LdBPK_366240.1; protein-l-isoaspartate o-methyltransferase (LdBPK_366860.1). |

### Table S7: Biological processes common between Host and *Leishmania*

| | Host | | | | *Leishmania* | | | |
|---|---|---|---|---|---|---|---|---|
| Name | Value | log_size | uniqueness | dispensability | Value | log_size | niqueness | spensabili |
| amide biosynthetic process | -12.926 | 5.552 | 0.963 | 0.229 | -18.781 | 5.552 | 0.888 | 0.229 |
| amide metabolic process | -12.718 | 5.87 | 0.983 | 0.064 | -19.342 | 5.87 | 0.961 | 0.064 |
| biosynthetic process | -13.297 | 7.064 | 0.974 | 0.104 | -11.751 | 7.064 | 0.941 | 0.104 |
| cell cycle | -3.439 | 5.921 | 0.989 | 0.021 | -1.875 | 5.921 | 0.979 | 0.021 |
| cell division | -2.655 | 5.748 | 0.99 | 0.02 | -2.626 | 5.748 | 0.98 | 0.02 |
| cellular biosynthetic process | -10.964 | 6.979 | 0.93 | 0.393 | -10.235 | 6.979 | 0.832 | 0.393 |
| cellular component biogenesis | -4.694 | 6.446 | 0.935 | 0.7 | -4.681 | 6.446 | 0.855 | 0.535 |
| cellular component organization | -7.745 | 6.7 | 0.93 | 0.693 | -2.801 | 6.7 | 0.844 | 0.7 |
| cellular component organization or biogenesis | -7.551 | 6.763 | 0.986 | 0.03 | -5.942 | 6.763 | 0.973 | 0.03 |
| cellular localization | -9.28 | 6.358 | 0.922 | 0.444 | -1.76 | 6.358 | 0.844 | 0.573 |
| cellular metabolic process | -12.863 | 7.207 | 0.963 | 0.209 | -4.225 | 7.207 | 0.918 | 0.277 |
| cellular process | -8.284 | 7.481 | 1 | 0 | -8.961 | 7.481 | 1 | 0 |
| cytoplasmic translation | -5.259 | 5.186 | 0.94 | 0.542 | -2.762 | 5.186 | 0.871 | 0.408 |
| energy coupled proton transport, down electrochemical gradient | -5.032 | 5.775 | 0.925 | 0.358 | -1.577 | 5.775 | 0.863 | 0.269 |
| ether lipid biosynthetic process | -2.621 | 3.523 | 0.932 | 0.385 | -1.468 | 3.523 | 0.828 | 0.44 |
| ether metabolic process | -1.921 | 4.192 | 0.957 | 0.256 | -1.468 | 4.192 | 0.903 | 0.041 |
| exocytosis | -3.478 | 5.153 | 0.915 | 0.527 | -1.304 | 5.153 | 0.877 | 0.659 |
| gene expression | -7.013 | 6.724 | 0.923 | 0.59 | -10.194 | 6.724 | 0.814 | 0.59 |
| localization | -6.293 | 6.902 | 1 | 0 | -1.75 | 6.902 | 1 | 0 |
| macromolecule biosynthetic process | -9.197 | 6.82 | 0.926 | 0.549 | -11.016 | 6.82 | 0.82 | 0.549 |
| macromolecule localization | -6.062 | 6.32 | 0.932 | 0.448 | -1.828 | 6.32 | 0.863 | 0.562 |
| macromolecule metabolic process | -7.192 | 7.132 | 0.974 | 0.225 | -4.902 | 7.132 | 0.939 | 0.193 |
| metabolic process | -13.246 | 7.373 | 1 | 0 | -6.142 | 7.373 | 1 | 0 |
| mitochondrial transport | -8.755 | 5.201 | 0.944 | 0.302 | -1.357 | 5.201 | 0.893 | 0.281 |
| nucleobase-containing compound transport | -2.093 | 5.367 | 0.929 | 0.521 | -2.087 | 5.367 | 0.861 | 0.502 |
| organelle organization | -11.77 | 6.425 | 0.93 | 0.626 | -2.324 | 6.425 | 0.849 | 0.618 |
| peptide biosynthetic process | -15.003 | 4.962 | 0.971 | 0.195 | -19.547 | 4.962 | 0.922 | 0.195 |
| peptide metabolic process | -14.548 | 5.21 | 0.985 | 0.053 | -18.868 | 5.21 | 0.967 | 0.053 |
| primary metabolic process | -9.258 | 7.296 | 0.972 | 0.277 | -4.739 | 7.296 | 0.934 | 0.25 |
| protein folding | -7.494 | 5.685 | 0.933 | 0.47 | -1.808 | 5.685 | 0.852 | 0.47 |
| protein import | -3.531 | 4.507 | 0.913 | 0.508 | -3.269 | 4.507 | 0.824 | 0.519 |
| protein localization to organelle | -5.156 | 5.737 | 0.901 | 0.691 | -2.367 | 5.737 | 0.784 | 0.603 |
| protein metabolic process | -6.62 | 6.885 | 0.957 | 0.306 | -8.864 | 6.885 | 0.903 | 0.306 |
| protein refolding | -7.626 | 4.898 | 0.942 | 0.697 | -1.343 | 4.898 | 0.874 | 0.697 |
| protein-containing complex organization | -4.867 | 6.063 | 0.936 | 0.488 | -2.052 | 6.063 | 0.863 | 0.455 |
| regulation of cell communication | -1.378 | 5.9 | 0.858 | 0.296 | -2.016 | 5.9 | 0.948 | 0.2 |
| regulation of cellular process | -1.704 | 7.004 | 0.827 | 0.598 | -1.324 | 7.004 | 0.934 | 0.374 |
| regulation of response to stimulus | -2.262 | 5.966 | 0.863 | 0.252 | -1.664 | 5.966 | 0.949 | 0.251 |
| regulation of signaling | -1.358 | 5.901 | 0.865 | 0.247 | -1.927 | 5.901 | 0.95 | 0.246 |
| response to abiotic stimulus | -3.637 | 5.578 | 0.946 | 0.347 | -1.558 | 5.578 | 0.98 | 0.284 |
| response to heat | -5.29 | 5.132 | 0.927 | 0.309 | -2.409 | 5.132 | 0.963 | 0 |
| translation | -15.379 | 6.255 | 0.928 | 0 | -19.641 | 6.255 | 0.833 | 0 |
| transport | -7.284 | 6.876 | 0.919 | 0.573 | -1.883 | 6.876 | 0.835 | 0.455 |

### Table S8: host vs host substrate

| | Host | | | | Host substrate | | | |
|---|---|---|---|---|---|---|---|---|
| Name | Value | log_size | uniqueness | dispensability | Value | log_size | uniqueness | dispensability |
| actin cytoskeleton organization | -5.481 | 5.499 | 0.928 | 0.577 | -4.939 | 5.499 | 0.909 | 0.018 |
| actin filament-based process | -4.949 | 5.514 | 0.99 | 0.019 | -5.186 | 5.514 | 0.991 | 0.018 |
| biological regulation | -1.76 | 7.056 | 1 | 0 | -7.864 | 7.056 | 1 | 0 |
| blood coagulation | -1.436 | 4.483 | 0.796 | 0.523 | -2.056 | 4.483 | 0.75 | 0.532 |
| canonical Wnt signaling pathway | -1.735 | 4.631 | 0.832 | 0.448 | -1.473 | 4.631 | 0.803 | 0.572 |
| cell death | -2.071 | 5.232 | 0.991 | 0.017 | -3.166 | 5.232 | 0.991 | 0.017 |
| cellular component assembly | -4.676 | 6.247 | 0.928 | 0.548 | -4.037 | 6.247 | 0.915 | 0.459 |
| cellular component biogenesis | -4.694 | 6.446 | 0.935 | 0.7 | -4.864 | 6.446 | 0.923 | 0.7 |
| cellular component organization | -7.745 | 6.7 | 0.93 | 0.693 | -5.369 | 6.7 | 0.918 | 0.693 |
| cellular component organization or biogenesis | -7.551 | 6.763 | 0.986 | 0.03 | -6.107 | 6.763 | 0.988 | 0.028 |
| cellular localization | -9.28 | 6.358 | 0.922 | 0.444 | -10.232 | 6.358 | 0.933 | 0.419 |
| cellular metabolic process | -12.863 | 7.207 | 0.963 | 0.209 | -2.115 | 7.207 | 0.976 | 0.225 |
| cellular process | -8.284 | 7.481 | 1 | 0 | -4.946 | 7.481 | 1 | 0 |
| cellular response to chemical stimulus | -7.505 | 5.946 | 0.906 | 0.022 | -2.29 | 5.946 | 0.903 | 0.618 |
| cellular response to chemical stress | -1.638 | 5.187 | 0.908 | 0.64 | -1.366 | 5.187 | 0.898 | 0.64 |
| cellular response to cytokine stimulus | -4.245 | 5.04 | 0.925 | 0.618 | -3.222 | 5.04 | 0.918 | 0.188 |
| cellular response to endogenous stimulus | -3.513 | 5.451 | 0.934 | 0.335 | -1.622 | 5.451 | 0.925 | 0.364 |
| cellular response to environmental stimulus | -1.984 | 4.813 | 0.942 | 0.332 | -5.44 | 4.813 | 0.932 | 0.357 |
| cellular response to external stimulus | -1.344 | 3.553 | 0.957 | 0.224 | -1.515 | 3.553 | 0.947 | 0.399 |
| cellular response to stimulus | -1.42 | 6.749 | 0.919 | 0.628 | -3.897 | 6.749 | 0.907 | 0.579 |
| cellular response to stress | -3.236 | 6.255 | 0.914 | 0.576 | -4.068 | 6.255 | 0.895 | 0.546 |
| cerebellum structural organization | -1.91 | 1.833 | 0.95 | 0.559 | -1.647 | 1.833 | 0.929 | 0.674 |
| coagulation | -1.426 | 4.484 | 0.957 | 0.41 | -2.039 | 4.484 | 0.936 | 0.469 |
| cytoplasmic translational initiation | -4.417 | 4.722 | 0.944 | 0.494 | -1.318 | 4.722 | 0.961 | 0.673 |
| cytoskeleton organization | -6.473 | 5.887 | 0.929 | 0.021 | -3.833 | 5.887 | 0.915 | 0.577 |
| developmental process | -4.548 | 6.192 | 1 | 0 | -2.505 | 6.192 | 1 | 0 |
| endomembrane system organization | -2.969 | 5.383 | 0.944 | 0.404 | -2.434 | 5.383 | 0.934 | 0.368 |
| establishment of Golgi localization | -1.316 | 2.403 | 0.93 | 0.565 | -2.465 | 2.403 | 0.933 | 0.565 |
| establishment of organelle localization | -4.316 | 5.294 | 0.926 | 0.302 | -4.777 | 5.294 | 0.931 | 0.309 |
| establishment or maintenance of cell polarity | -4.444 | 4.964 | 0.991 | 0.016 | -2.629 | 4.964 | 0.992 | 0.016 |
| export from cell | -1.926 | 5.56 | 0.933 | 0.332 | -1.643 | 5.56 | 0.943 | 0.333 |
| gene expression | -7.013 | 6.724 | 0.923 | 0.59 | -4.004 | 6.724 | 0.948 | 0.455 |
| isotype switching to IgG isotypes | -2.21 | 1.415 | 0.877 | 0.653 | -1.945 | 1.415 | 0.855 | 0.683 |
| localization | -6.293 | 6.902 | 1 | 0 | -5.214 | 6.902 | 1 | 0 |
| mRNA metabolic process | -2.211 | 5.83 | 0.941 | 0.31 | -6.949 | 5.83 | 0.951 | 0.528 |
| mRNA modification | -1.986 | 4.458 | 0.951 | 0.407 | -1.858 | 4.458 | 0.954 | 0.69 |
| macromolecule localization | -6.062 | 6.32 | 0.932 | 0.448 | -3.377 | 6.32 | 0.942 | 0.448 |
| macromolecule metabolic process | -7.192 | 7.132 | 0.974 | 0.225 | -2.394 | 7.132 | 0.987 | 0.163 |
| membrane organization | -2.2 | 5.683 | 0.941 | 0.437 | -3.584 | 5.683 | 0.93 | 0.485 |
| metabolic process | -13.246 | 7.373 | 1 | 0 | -3.935 | 7.373 | 1 | 0 |
| midbrain development | -1.52 | 3.254 | 0.943 | 0.653 | -1.73 | 3.254 | 0.92 | 0.451 |
| mitochondrial membrane organization | -4.365 | 4.753 | 0.936 | 0.673 | -1.591 | 4.753 | 0.928 | 0.536 |
| mitochondrial transport | -8.755 | 5.201 | 0.944 | 0.302 | -1.657 | 5.201 | 0.953 | 0.302 |
| mitotic cell cycle | -2.216 | 5.645 | 0.977 | 0.448 | -1.738 | 5.645 | 0.971 | 0.019 |
| negative regulation of amide metabolic process | -2.387 | 2.9 | 0.871 | 0.418 | -1.562 | 2.9 | 0.83 | 0.418 |
| negative regulation of biological process | -5.245 | 6.278 | 0.854 | 0.312 | -8.935 | 6.278 | 0.819 | 0.304 |
| negative regulation of cellular catabolic process | -1.857 | 4.327 | 0.829 | 0.698 | -2.015 | 4.327 | 0.779 | 0.698 |
| negative regulation of cellular process | -4.905 | 6.243 | 0.801 | 0.289 | -9.51 | 6.243 | 0.737 | 0.325 |
| negative regulation of integrin-mediated signaling pathway | -1.735 | 2.814 | 0.862 | 0.43 | -1.473 | 2.814 | 0.819 | 0.47 |
| nitrogen compound transport | -7.771 | 6.326 | 0.929 | 0.579 | -2.79 | 6.326 | 0.94 | 0.579 |
| nucleic acid metabolic process | -1.482 | 6.736 | 0.929 | 0.644 | -6.171 | 6.736 | 0.955 | 0.415 |
| nucleobase-containing compound metabolic process | -5.614 | 6.873 | 0.967 | 0.154 | -4.366 | 6.873 | 0.984 | 0.117 |
| nucleobase-containing compound transport | -2.093 | 5.367 | 0.929 | 0.521 | -1.63 | 5.367 | 0.943 | 0.502 |
| organelle localization | -4.494 | 5.469 | 0.943 | 0.308 | -5.679 | 5.469 | 0.952 | 0.316 |
| organelle organization | -11.77 | 6.425 | 0.93 | 0.626 | -6.364 | 6.425 | 0.917 | 0.626 |
| positive regulation of biological process | -3.593 | 6.254 | 0.855 | 0.313 | -8.795 | 6.254 | 0.82 | 0.313 |
| positive regulation of cellular process | -3.984 | 6.174 | 0.775 | 0.325 | -10.499 | 6.174 | 0.75 | 0 |
| post-transcriptional regulation of gene expression | -5.3 | 5.822 | 0.839 | 0.251 | -3.478 | 5.822 | 0.794 | 0.335 |
| primary metabolic process | -9.258 | 7.296 | 0.972 | 0.277 | -1.851 | 7.296 | 0.987 | 0.277 |
| protein refolding | -7.626 | 4.898 | 0.942 | 0.697 | -1.714 | 4.898 | 0.96 | 0.326 |
| protein-containing complex organization | -4.867 | 6.063 | 0.936 | 0.488 | -3.029 | 6.063 | 0.924 | 0.548 |
| regulation of actin filament-based process | -2.439 | 5.139 | 0.876 | 0.185 | -3.216 | 5.139 | 0.849 | 0.233 |
| regulation of amide metabolic process | -4.981 | 4.034 | 0.886 | 0.252 | -2.939 | 4.034 | 0.857 | 0.227 |
| regulation of anatomical structure size | -3.447 | 5.196 | 0.86 | 0.599 | -2.427 | 5.196 | 0.84 | 0.483 |
| regulation of apoptotic process | -3.842 | 5.301 | 0.854 | 0.22 | -4.043 | 5.301 | 0.836 | 0.243 |
| regulation of binding | -3.184 | 3.334 | 0.881 | 0.583 | -2.011 | 3.334 | 0.872 | 0.568 |
| regulation of biological process | -1.52 | 7.035 | 0.833 | 0.676 | -8.376 | 7.035 | 0.789 | 0.676 |
| regulation of biological quality | -5.348 | 6.09 | 0.866 | 0.238 | -2.355 | 6.09 | 0.835 | 0.27 |
| regulation of catabolic process | -3.027 | 5.523 | 0.855 | 0.349 | -2.004 | 5.523 | 0.816 | 0.349 |
| regulation of cell communication | -1.378 | 5.9 | 0.858 | 0.296 | -3.829 | 5.9 | 0.825 | 0.29 |
| regulation of cell-substrate junction organization | -3.18 | 3.688 | 0.836 | 0.558 | -1.344 | 3.688 | 0.83 | 0.558 |
| regulation of cellular catabolic process | -2.771 | 4.961 | 0.85 | 0.32 | -2.863 | 4.961 | 0.816 | 0.269 |
| regulation of cellular component biogenesis | -5.954 | 5.358 | 0.872 | 0.223 | -2.921 | 5.358 | 0.843 | 0.247 |
| regulation of cellular component organization | -7.697 | 5.773 | 0.862 | 0.217 | -5.91 | 5.773 | 0.83 | 0.278 |
| regulation of cellular localization | -5.966 | 5.029 | 0.815 | 0.18 | -4.018 | 5.029 | 0.814 | 0.519 |
| regulation of cellular process | -1.704 | 7.004 | 0.827 | 0.598 | -9.362 | 7.004 | 0.781 | 0.435 |
| regulation of developmental process | -1.72 | 5.764 | 0.868 | 0.229 | -4.387 | 5.764 | 0.838 | 0.257 |
| regulation of growth | -2.564 | 4.892 | 0.887 | 0.175 | -2.67 | 4.892 | 0.862 | 0.203 |
| regulation of localization | -4.229 | 5.566 | 0.873 | 0.205 | -5.836 | 5.566 | 0.845 | 0.242 |
| regulation of locomotion | -2.389 | 5.219 | 0.881 | 0.189 | -3.325 | 5.219 | 0.854 | 0.22 |
| regulation of mast cell apoptotic process | -1.516 | 1.792 | 0.883 | 0.561 | -1.473 | 1.792 | 0.882 | 0.561 |
| regulation of membrane permeability | -2.066 | 3.918 | 0.884 | 0.475 | -2.462 | 3.918 | 0.869 | 0.156 |
| regulation of metabolic process | -2.576 | 6.806 | 0.835 | 0.394 | -6.737 | 6.806 | 0.793 | 0.598 |
| regulation of molecular function | -8.18 | 5.771 | 0.874 | 0 | -7.306 | 5.771 | 0.846 | 0.244 |
| regulation of multicellular organismal process | -1.425 | 5.543 | 0.874 | 0.204 | -2.536 | 5.543 | 0.845 | 0.24 |
| regulation of programmed cell death | -4.156 | 5.315 | 0.873 | 0.22 | -3.895 | 5.315 | 0.844 | 0.244 |
| regulation of protein metabolic process | -5.118 | 6.003 | 0.839 | 0.41 | -3.357 | 6.003 | 0.793 | 0.41 |
| regulation of protein-containing complex assembly | -6.566 | 5.049 | 0.792 | 0.181 | -5.141 | 5.049 | 0.786 | 0.228 |
| regulation of response to stimulus | -2.262 | 5.966 | 0.863 | 0.252 | -5.265 | 5.966 | 0.831 | 0.273 |
| regulation of signaling | -1.358 | 5.901 | 0.865 | 0.247 | -4.526 | 5.901 | 0.834 | 0.268 |
| response to abiotic stimulus | -3.637 | 5.578 | 0.946 | 0.347 | -2.385 | 5.578 | 0.935 | 0.378 |
| response to endogenous stimulus | -5.004 | 5.525 | 0.946 | 0.342 | -1.911 | 5.525 | 0.935 | 0.372 |
| response to gamma radiation | -2.237 | 3.387 | 0.948 | 0.601 | -3.332 | 3.387 | 0.937 | 0.654 |
| response to methamphetamine hydrochloride | -1.91 | 1.114 | 0.951 | 0.43 | -1.647 | 1.114 | 0.955 | 0.287 |
| response to oxidative stress | -2.559 | 5.494 | 0.937 | 0.636 | -1.469 | 5.494 | 0.919 | 0.636 |
| response to stimulus | -1.993 | 6.838 | 1 | 0 | -2.243 | 6.838 | 1 | 0 |
| response to stress | -4.635 | 6.428 | 0.935 | 0.503 | -3.532 | 6.428 | 0.922 | 0.628 |
| ruffle organization | -1.862 | 3.246 | 0.959 | 0.448 | -1.428 | 3.246 | 0.949 | 0.427 |
| secretion | -2.044 | 5.504 | 0.941 | 0.327 | -1.65 | 5.504 | 0.951 | 0.327 |
| supramolecular fiber organization | -4.352 | 5.506 | 0.943 | 0.417 | -3.709 | 5.506 | 0.932 | 0.46 |
| synaptic target recognition | -2.21 | 2.531 | 0.944 | 0.415 | -1.945 | 2.531 | 0.923 | 0.485 |
| transport | -7.284 | 6.876 | 0.919 | 0.573 | -4.564 | 6.876 | 0.932 | 0.573 |
| vesicle organization | -2.512 | 5.297 | 0.937 | 0.548 | -1.57 | 5.297 | 0.924 | 0.5 |
| vesicle-mediated transport | -4.566 | 6.022 | 0.926 | 0.381 | -4.501 | 6.022 | 0.937 | 0.383 |

Table S9: Biological processes only *Leishmania*

| description | dispensability | log_size | uniqueness | value |
|---|---|---|---|---|
| **cellular response to amino acid starvation** | 0.384 | 4.341 | 0.957 | -1.468 |
| **regulation of TOR signaling** | 0.117 | 4.757 | 0.898 | -2.904 |
| **positive regulation of TORC1 signaling** | 0.64 | 4.234 | 0.892 | -2.766 |
| **regulation of protein acetylation** | 0.091 | 2.778 | 0.958 | -1.766 |
| **cellular response to antimycin A** | 0.127 | 0.301 | 0.978 | -1.766 |
| **response to antimycin A** | 0.356 | 0.301 | 0.986 | -1.766 |
| **non-membrane-bounded organelle assembly** | 0.592 | 5.591 | 0.836 | -5.172 |
| **membrane invagination** | 0.575 | 4.299 | 0.903 | -1.838 |
| **plasma membrane invagination** | 0.292 | 4.293 | 0.903 | -1.838 |
| **exocyst localization** | 0.176 | 3.745 | 0.921 | -1.468 |
| **organelle assembly** | 0.682 | 5.804 | 0.835 | -2.519 |
| **regulation of peroxisome size** | 0.348 | 3.097 | 0.889 | -1.468 |
| **regulation of membrane potential** | 0.463 | 5.198 | 0.951 | -2.766 |
| **regulation of mitochondrial membrane potential** | 0 | 3.657 | 0.961 | -3.062 |
| **cytokinesis** | 0.017 | 5.129 | 0.929 | -2.848 |
| **mitotic cytokinetic process** | 0.605 | 4.02 | 0.935 | -1.838 |
| **mitotic cell cycle phase transition** | 0.642 | 4.788 | 0.924 | -1.764 |
| **cell cycle phase transition** | 0.642 | 4.792 | 0.934 | -1.696 |
| **carboxylic acid biosynthetic process** | 0.499 | 6.234 | 0.779 | -1.334 |
| **thioester metabolic process** | 0.133 | 5.306 | 0.906 | -1.696 |
| **oligosaccharide biosynthetic process** | 0.188 | 4.817 | 0.884 | -1.468 |
| **isoprenoid biosynthetic process** | 0.263 | 5.295 | 0.846 | -1.696 |
| **isoprenoid metabolic process** | 0.569 | 5.381 | 0.885 | -1.523 |
| **proline biosynthetic process** | 0.607 | 4.65 | 0.837 | -1.468 |
| **L-leucine metabolic process** | 0.233 | 4.959 | 0.852 | -1.468 |
| **fructose 6-phosphate metabolic process** | 0.112 | 4.65 | 0.874 | -2.766 |
| **nucleoside triphosphate metabolic process** | 0.645 | 5.675 | 0.783 | -1.697 |
| **purine ribonucleoside triphosphate metabolic process** | 0.435 | 5.555 | 0.755 | -2.342 |
| **purine ribonucleotide metabolic process** | 0.69 | 5.819 | 0.766 | -1.379 |
| **ribonucleoprotein complex biogenesis** | 0.673 | 6.036 | 0.839 | -8.2 |
| **post-transcriptional tethering of RNA polymerase II gene DNA at nuclear periphery** | 0.505 | 3.917 | 0.903 | -2.766 |
| **transcription-dependent tethering of RNA polymerase II gene DNA at nuclear periphery** | 0.288 | 4.226 | 0.898 | -4.021 |
| **maturation of LSU-rRNA** | 0.695 | 4.846 | 0.725 | -2.687 |
| **ribosome biogenesis** | 0.022 | 5.965 | 0.82 | -8.816 |
| **protein import into nucleus** | 0.016 | 5.004 | 0.769 | -4.539 |
| **protein-containing complex localization** | 0.219 | 5.035 | 0.9 | -3.534 |
